# Carbohydrate and PepO control bimodality in competence development by *Streptococcus mutans*

**DOI:** 10.1101/670752

**Authors:** Simon A.M. Underhill, Robert C. Shields, Robert A. Burne, Stephen J. Hagen

## Abstract

In *Streptococcus mutans*, the alternative sigma factor ComX controls entry into genetic competence. Competence signaling peptide (CSP) induces bimodal expression of *comX*, with only a fraction of cells in the population becoming transformable. Curiously, bimodal *comX* activation in response to CSP is affected by peptides in the growth medium and by carbohydrate source. CSP elicits bimodal expression of *comX* in media rich in small peptides, but in defined media lacking small peptides CSP induces no response in *comX*. In addition, growth on certain sugars other than glucose increases the proportion of the population that activates *comX* in response to CSP, relative to growth on glucose. By investigating the connection between media and bimodal *comX* expression, we find evidence for two mechanisms that modulate transcriptional positive feedback in the ComRS system, which is the origin of *comX* bimodality. We find that the endopeptidase PepO suppresses the ComRS feedback loop, most likely by degrading the intracellular XIP/ComS signal. Deletion of *pepO* eliminates bimodality in *comX*, leading to a unimodal *comX* response to CSP in defined and complex media. We also find that CSP upregulates *comR* in a carbohydrate source-dependent fashion, providing an additional stimulus to the ComRS feedback system. Our data provide mechanistic insight into how CSP regulates the bistable competence circuit and explain the puzzle of growth medium-dependence in *S. mutans* competence regulation.

## Introduction

The Gram-positive bacterium *Streptococcus mutans* inhabits human oral biofilms and is a primary etiologic agent of dental caries (Loesche, 1986). The ability of *S. mutans* to colonize the oral cavity and compete with commensal organisms is associated with the *com* regulon. The *com* (competence) regulon controls entry into genetic competence, a transient state during which cells are able to internalize DNA from the environment (Shanker; Federle, 2017). In *S. mutans* the *com* regulon is linked to tolerance of environmental stress, including heat, oxidative stresses, pH and carbohydrate availability (Qi *et al.*, 2004; Ahn *et al.*, 2005; Ahn *et al.*, 2006; Senadheera, M. D. *et al.*, 2007; Tremblay *et al.*, 2009; Senadheera, D. B. *et al.*, 2012). It is also involved in bacteriocin production and lysis, which are important in interspecies competition (Shanker; Federle, 2017), and in biofilm formation and stability (Li *et al.*, 2002). There are many unresolved questions in the study of *S. mutans* competence, especially in regard to how the *com* regulon integrates diverse environmental cues, such as the presence of different signal peptides or nutrients or the composition of the growth medium, in order to drive different phenotypic outputs in downstream regulated genes (Hagen; Son, 2017).

The quorum sensing peptide CSP (<u>c</u>ompetence <u>s</u>timulating <u>p</u>eptide) (Li *et al.*, 2002) is a primary input to the *S. mutans* competence pathway (Fig. 1). CSP is derived from a 46-residue precursor encoded by *comC*, processed to 21 residue length, then exported to the extracellular environment by the ComAB transporter. The ComC peptide is further processed by the extracellular SepM protease to yield the mature 18-residue CSP, which is understood to be the most active form of the peptide (Hossain; Biswas, 2012; Shanker; Federle, 2017). Extracellular CSP stimulates the competence pathway by interacting with the ComDE two-component signal transduction system (TCSTS): CSP bound by the transmembrane kinase ComD induces phosphorylation of ComE, which then acts as a transcriptional activator for several bacteriocin and competence-related genes, including *cipB* (Perry *et al.*, 2009; Fontaine *et al.*, 2015). Although the mechanism is not known, expression of *cipB* stimulates the ComRS system, an Rgg-type signaling system that is the immediate regulator of *comX* (also called *sigX*). ComX (or SigX) is an alternative sigma factor for late competence genes, which encode proteins for uptake and processing of DNA. Therefore, CSP drives transformability through a pathway that includes ComCDE, *cipB*, ComRS and ComX, with the regulatory link from *cipB* to ComRS being the least understood. However, a signature characteristic of this pathway (unlike the generally similar pathway in *S. pneumoniae* (Shanker; Federle, 2017)) is that *comX* responds bimodally (if at all) to CSP: when CSP is supplied in complex media containing small peptides, *comX* activates in only a subpopulation of cells, even though *cipB* activates population-wide. In defined media lacking small peptides, CSP elicits no *comX* response, although *cipB* is again activated population wide (Son, Minjun *et al.*, 2015; Reck *et al.*, 2015).

**Fig. 1:**
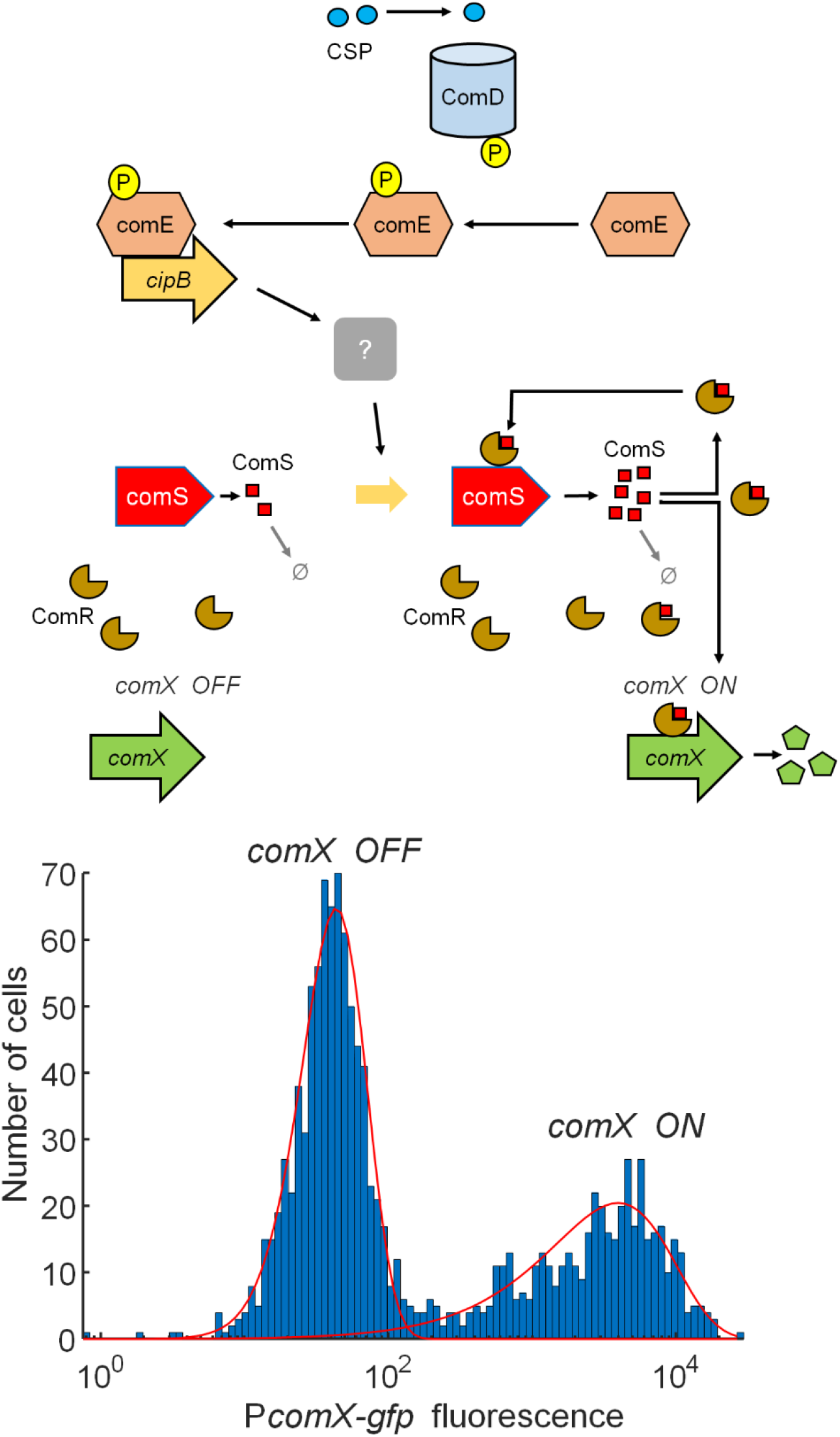
current model of CSP induction of competence. CSP induction of competence in *S. mutans* (Shanker; Federle, 2017; Underhill *et al.*, 2018). CSP binds the ComD transmembrane kinase, which phosphorylates ComE. ComE phosphate activates bacteriocin genes including *cipB*, which through an unknown mechanism stimulates the ComRS positive feedback system. ComS interacts with ComR to form a transcriptional activator for *comS* and *comX*. Positive transcriptional feedback in *comS* amplifies fluctuations in ComS levels, allowing cells to stochastically flip the ComRS system from a transcriptionally *OFF* to *ON* state and drive *comX* transcription. Degradation of basal ComS by an unidentified peptidase is hypothesized to inhibit activation of ComRS (Son, M. *et al.*, 2012). Cell to cell variability in the state of the ComRS switch leads to bimodal expression of a P*comX-gfp* reporter in a population of *S. mutans*. The inset at bottom shows a histogram of individual cell reporter fluorescence for cells supplied 1 μM CSP in complex growth medium (TV). Overlaid (red curves) are gamma probability distribution functions corresponding to OFF (left) and ON (right) states of the ComRS/*comX* system. (*Methods*).

Bimodality arises in the ComRS system, which lies downstream of *cipB* (Son, M. *et al.*, 2012). Mashburn-Warren *et al.* (Mashburn-Warren *et al.*, 2010) showed that XIP (*sig<u>X</u>*-<u>i</u>nducing <u>p</u>eptide), comprising the C-terminal 7 residues of ComS, can induce *comX* if supplied exogenously to *S. mutans*. XIP is imported by the Opp permease and interacts with ComR to form a multimeric complex that is a transcriptional activator for both *comX* and *comS* (Mashburn-Warren *et al.*, 2010; Son, M. *et al.*, 2012; Fontaine *et al.*, 2013; Underhill *et al.*, 2018). XIP induces a population-wide (unimodal) *comX* response when provided as an extracellular signal in defined growth medium. However the presence of transcriptional positive feedback via *comS* also allows the ComRS system to operate as an intracellular positive feedback loop: endogenously produced ComS apparently interacts with ComR intracellularly to enhance transcription of *comS* and *comX* (Underhill *et al.*, 2018). Such a feedback mechanism is sensitive to basal levels of ComR and ComS, which vary stochastically among cells (Dubnau; Losick, 2006). Consequently the positive feedback behavior is heterogeneous and only a subpopulation flips the ComRS switch by expressing *comX* and *comS* above basal levels, ultimately resulting in a bimodal distribution of ComX in the population (Fig. 1).

How CSP activates this feedback loop remains unclear. The pathway through which *cipB* (or other upstream elements) stimulates ComRS has not been identified. In addition, it is not understood why CSP elicits a bimodal *comX* response only in complex growth medium that contains small peptides. A related question is why CSP elicits no *comX* response in defined media that lacks small peptides. We have suggested that small peptides imported from the growth medium could strengthen ComRS feedback – and thus favor *comX* activation – by competing for an *S. mutans* enzyme that degrades endogenously produced ComS (Son, M. *et al.*, 2012; Hagen; Son, 2017). However, no such enzyme has yet been identified. An additional question is what limits the proportion of cells that activate *comX* in the presence of CSP. Although the number of *comX*-active cells initially increases as the concentration of exogenous CSP is increased, the proportion of cells that activate *comX* expression saturates at an upper limit of 30-35% (Son, M. *et al.*, 2012).

A previous study (Moye *et al.*, 2016) identified carbon source as one of the few parameters – other than complex/defined medium - that alters the proportion of cells activating the ComRS switch, i.e. the probability of transition from *comX* “OFF” to “ON”. Single-cell studies found that CSP induced higher ON fractions in fructose- or trehalose-grown cultures than in glucose-grown cultures (Moye *et al.*, 2016), with trehalose giving the strongest response. In addition, CSP-stimulated cells growing in maltose or sucrose expressed a P*comX-lacZ* (*comX* promoter fusion) reporter at higher levels than did glucose-grown cells. In addition, growth on trehalose or sucrose led to higher transformation efficiencies than did growth on glucose. Deletion of the gene for the global carbon catabolite repression (CCR) mediator CcpA eliminated differences in CSP response between sugars (Moye *et al.*, 2016). Moye *et. al.* (Moye *et al.*, 2016) did not rule out the possibility that carbohydrate influences *comDE* or *comC* expression. However, the fact that *cipB* expression already saturates at moderate CSP levels, and that higher levels of CSP do not increase the fraction of responding cells (Son, M. *et al.*, 2012), suggests that the enhanced *comX* response in media formulated with carbohydrates other than glucose is not due to upregulation of *comCDE*.

Given the link between bimodality and carbohydrate source (Moye *et al.*, 2016), and our hypothesis that a peptidase could govern bimodality by modulating the strength of feedback in the ComRS system (Hagen; Son, 2017), we investigated whether *S. mutans* PepO – whose homologs are known to interact with Rgg signaling in other streptococci (Wilkening *et al.*, 2016) – could mediate the carbohydrate effect in *S. mutans* by interfering with ComRS autoactivation. We studied the relationship between carbohydrate source and activation of the CSP-induced competence pathway, using fluorescent gene reporter studies of individual cells, and biochemical and transcriptional approaches. By showing a clear connection between *pepO* and bimodality in ComRS, and demonstrating carbohydrate-sensitive effects on ComR regulation, our studies identify two mechanisms by which *S. mutans* controls the proportion of cells that enter the competent state.

## Results

### Trehalose enhances *comX*, but not *cipB*, activation by CSP

To confirm a previous report (Moye *et al.*, 2016) that growth in trehalose leads to a greater proportion of *comX*-ON cells in response to CSP than does growth in glucose, we measured the proportion of *comX*-ON cells as a function of the glucose and trehalose content of the growth medium. We grew a P*comX-gfp* reporter strain of *S. mutans* in complex growth medium (TV, *Methods*) containing mixtures of glucose and trehalose. Ratios of glucose to trehalose were chosen to maintain a constant hexose concentration of 20 mM, such that 2[tre]+[glc] = 20 mM (Moye *et al.*, 2016). We added 1 μM CSP-18 to each sample as it reached OD_600_ 0.1. The fluorescent reporter activity of individual cells was measured and histogrammed as described in (Kwak *et al.*, 2012). Growth of the two strains was checked to ensure these effects were not due to poor growth of the mutant on trehalose admixtures; the Δ*treR* strain grows poorly on trehalose but normally in the mixed carbohydrate media (Supplemental Figure S1). Fig. 2A shows that the presence of 0.5 to 1 mM trehalose was sufficient to increase the proportion of *comX*-ON cells. Repeating the experiment using the same *comX* reporter in a Δ*treR* strain showed no effect of trehalose on the proportion of *comX*-ON cells. As a Δ*treR* strain lacks the TreR transcriptional activator and does not express *treAB* (Baker *et al.*, 2018), these findings confirm that the trehalose-induced increase in the proportion of cells responding to CSP requires activation of the *tre* operon.

**Fig. 2:**
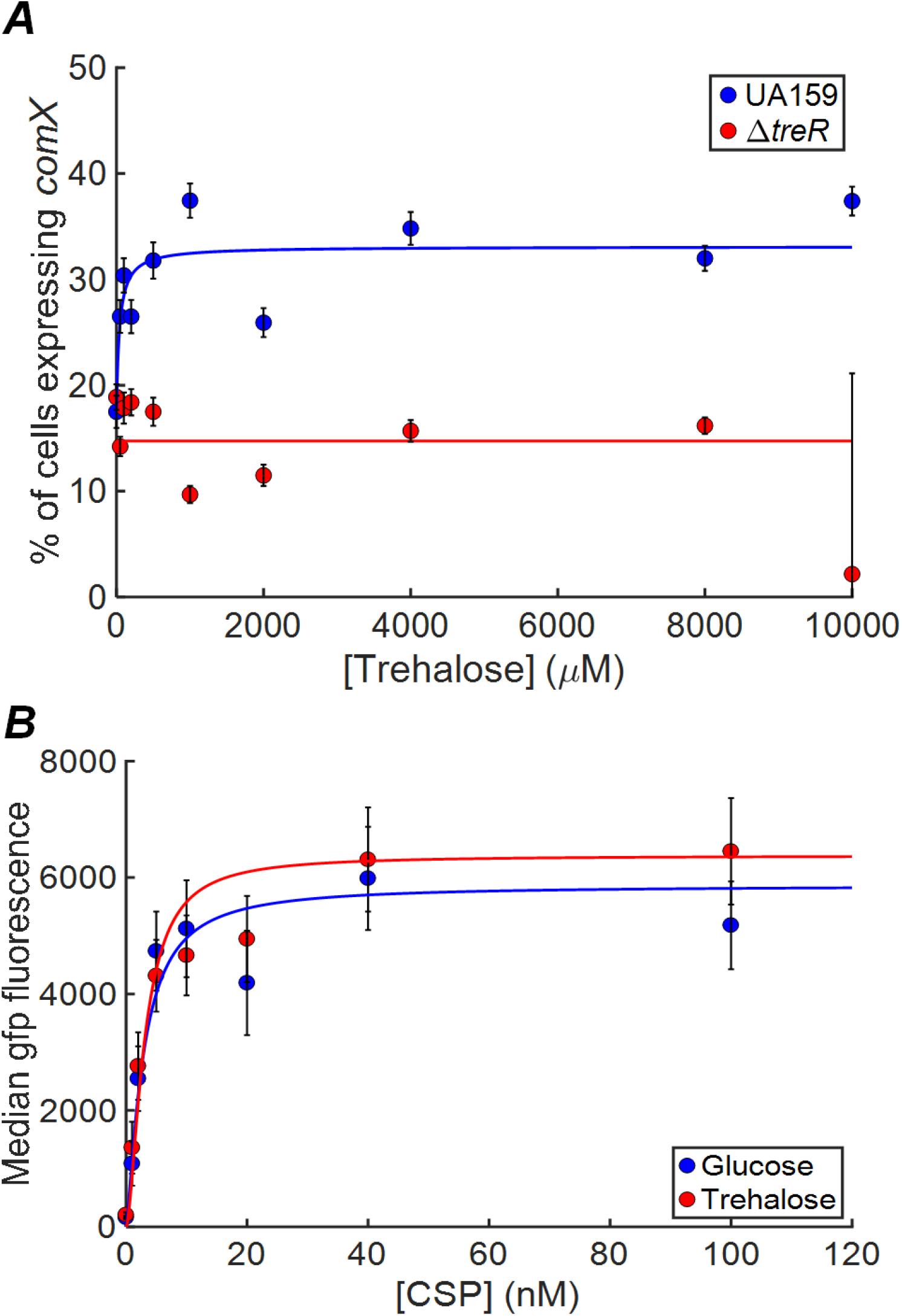
replacement of glucose by trehalose produces graded increase in percentage of cells responding to CSP. (A) Effect on *comX* activation of replacing glucose with trehalose as the carbon source. Carbohydrate composition of the TV medium was adjusted subject to the constraint 2[tre] + [glc] = 20 mM. In each sample 1 μM CSP was added at OD_600_ 0.1, and cell responses were measured by fluorescence microscopy. Data show the percentage of cells expressing *comX* vs. [CSP] in the wild type reporting background (blue) and a *treR* mutant (mutant that does not express the *tre* operon, red). Solid curves of corresponding colors represent *n* = 1 Hill function fits to the curves. (B) Median fluorescence of P*cipB-gfp* activity in glucose- (blue) and trehalose- (red) grown cells.

The activation of *cipB* transcription by phosphorylated ComE is an early step in the CSP-induced competence pathway (Son, Minjun *et al.*, 2015). We next tested whether alterations in *cipB* regulation, required for CSP stimulation (Perry *et al.*, 2009), could be triggered by carbohydrate. In order to test whether carbon source affects the competence pathway by influencing *cipB* or *comCDE* activity, we studied the effect of glucose/trehalose on the CSP response of a P*cipB-gfp* reporter strain. Fig. 2B shows that induction of *cipB* transcription by CSP was not significantly different in cells growing on trehalose compared to glucose. Fitting the median (in the cell population) *cipB* expression level to a simple binding isotherm (Hill function with *n* = 1), we find indistinguishable constants *K* = 2.6 ± 1.4 nM for glucose and 3.0 ± 0.9 nM for trehalose. These data indicate that *cipB* is not differentially regulated in trehalose versus glucose. The carbohydrate effect on *comX* expression must arise elsewhere than in the CSP-ComDE circuit.

### The carbohydrate effect on transformation efficiency is CSP-dependent

Because *S. mutans* maintains a low level of transformability even in the absence of CSP (Perry *et al.*, 2009), it was necessary to test whether the carbohydrate effect on transformability occurs through the CSP-induction pathway. We compared the transformation efficiency of wild-type UA159, a *pepO* mutant and a *comDE* mutant, grown in TV that contained mixtures of glucose and trehalose, with or without CSP. Fig. 3 shows that in the absence of CSP, the efficiency of transformation of wild-type cells is only slightly lower in glucose than in trehalose (*P* = 0.134 by Student’s *t* test). When CSP is provided, the transformation efficiency of the wild type is significantly greater in trehalose than in glucose (*P* < 0.001). These data imply that the carbohydrate effect is facilitated, if not entirely generated, by the CSP induction pathway. Consistent with this interpretation, we find that the CSP dependent differences in transformability are absent in the *comDE* deletion, as are significant differences between the different carbohydrates.

**Fig. 3:**
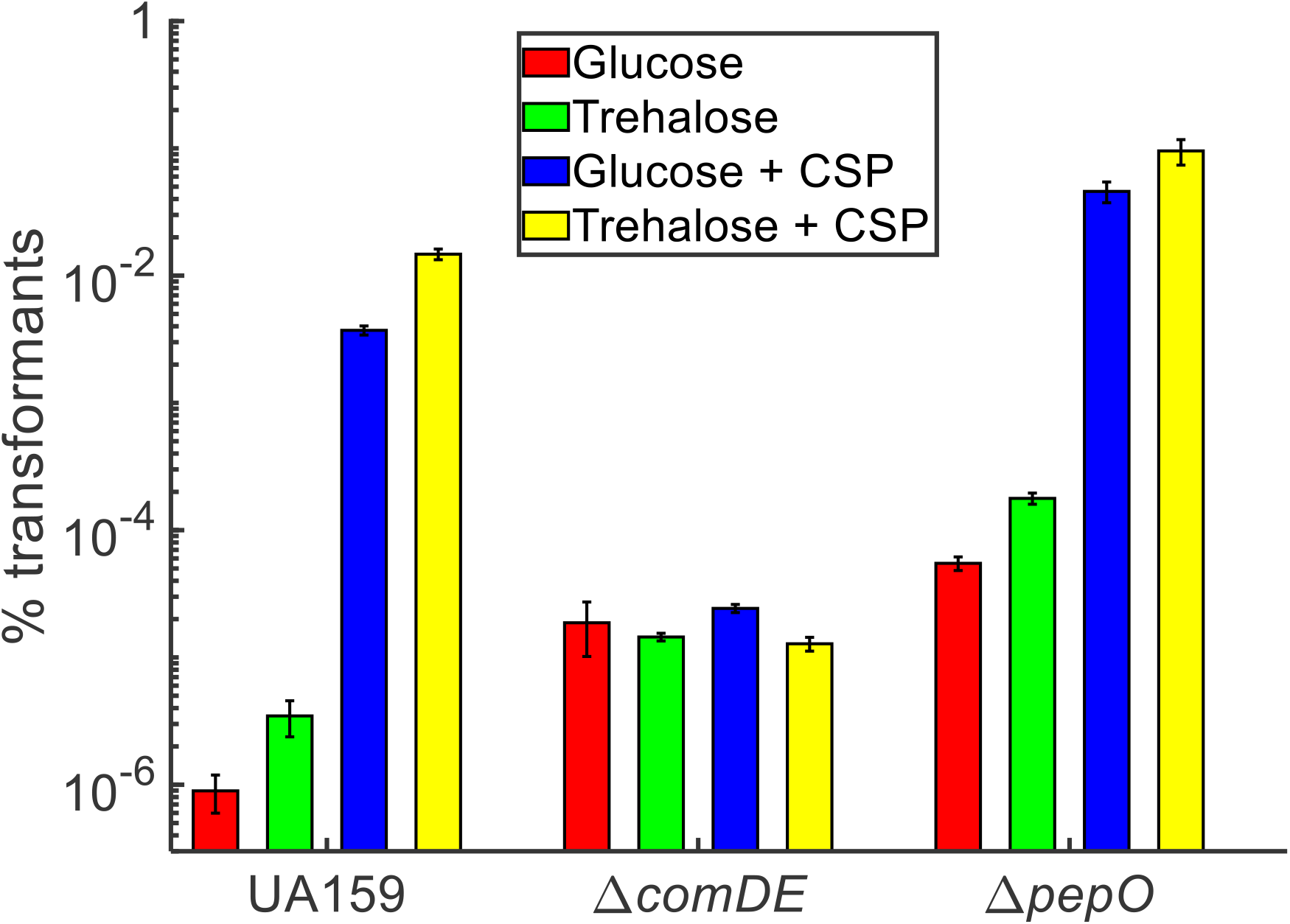
transformation efficiency is affected by carbohydrate only in the presence of CSP. Transformation efficiency (expressed as % transforming) in indicated genetic backgrounds in glucose (red), trehalose (green), glucose with 1 μM added CSP (blue) or trehalose with 1 μM added CSP (yellow).

Fig. 3 also shows that deletion of *pepO* increases the transformation efficiency under all conditions, both in the presence and absence of CSP. In the absence of CSP the Δ*pepO* strain had a roughly 100-fold higher rate of transformation than the wild type in glucose (*P* < 0.001) and in trehalose (*P* = 0.01). With CSP the difference narrowed to 30-fold in glucose (*P* = 0.038) and 16-fold in trehalose (*P* = 0.064). The Δ*pepO* strain exhibited a higher statistical significance in the carbohydrate effect (glucose vs. trehalose) of the non-CSP transformability (*P* = 0.012) than UA159. Therefore, deletion of *pepO* generally enhances transformability, but it does not eliminate CSP-sensitivity or the effect of carbohydrate on transformability.

### Carbohydrate-dependent increases in *comR* transcription in response to CSP

In order to investigate how certain carbohydrates could affect the ComRS bimodal switch compared to glucose, we used RT-qPCR to compare *comR* transcript levels (normalized to 16S rRNA) in cells grown in TV supplemented with glucose, trehalose or maltose, in the presence of 0 nM, 4 nM or 400 nM CSP. Fig. 4A shows that *comS* transcription was generally elevated in the presence of CSP, as expected if CSP drives competence by promoting *comS* transcriptional feedback (Hagen; Son, 2017; Underhill *et al.*, 2018). The CSP enhancement of *comS* transcription was greater in the tested sugars than in glucose and was greater in the *pepO* deletion. Less expected was the finding (Fig. 4B) that addition of CSP to the wild-type strain in trehalose or maltose caused a modest but significant 2-3 fold increase (*P* < 0.001) in *comR* transcripts, which did not occur in glucose. A CSP enhancement of *comR* transcription was also seen in the Δ*pepO* strain in those carbohydrates. We note that Lemme *et al.* (Lemme *et al.*, 2011) reported a similar, 1.4-1.8-fold upregulation of *comR* in cells that were treated with CSP in Todd-Hewitt/yeast extract medium. Curiously, we found no significant increase in *comR* transcripts when CSP was provided to wild-type cells growing in glucose. The finding that CSP leads to a modest upregulation of *comR* transcription, especially in the disaccharides tested here, suggests a mechanism by which CSP could stimulate the ComRS system and promote *comX* activity. As the bistable behavior of the ComRS feedback system will be sensitive to basal levels of both ComS and ComR, even a modest upregulation of *comR* by CSP, as occurs in trehalose and maltose, would promote positive feedback in the ComRS circuit, permitting a larger proportion of cells to enter the *comX*-ON state.

**Fig. 4:**
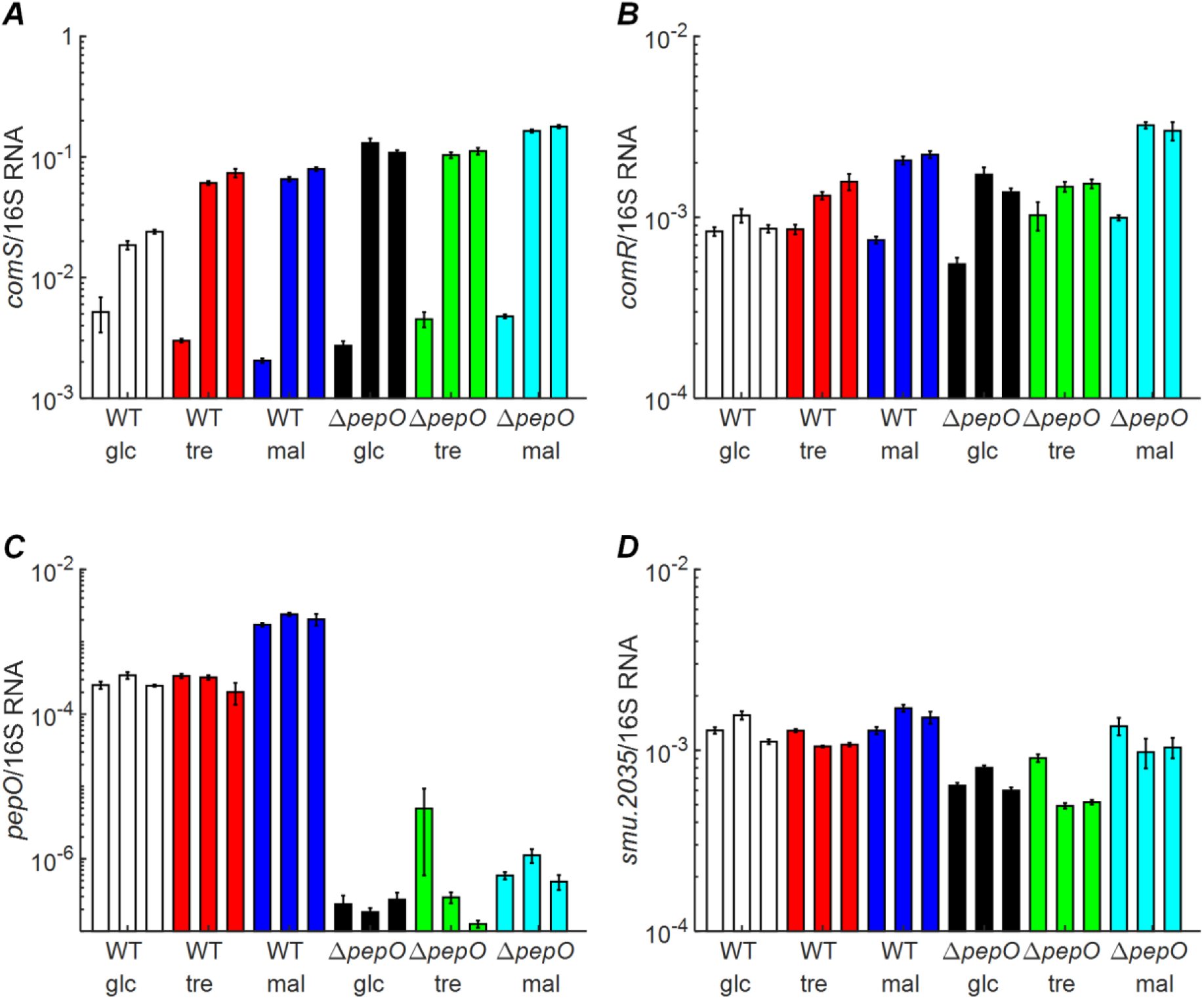
RT-qPCR measurement of gene expression modulation by sugar and CSP. RT-qPCR measurements of indicated transcripts normalized to 16S rRNA. Bars are grouped by strain and carbohydrate in color and correspond to 0, 4 nM and 400 nM added CSP from left to right within a color group. (A) *comS* expression. (B) *comR* expression. (C) *pepO* expression. (D) *smu.2035* expression.

Although our transformation assay (Fig. 3) demonstrated that deletion of *pepO* enhances *com* activity, Fig. 4C shows that *pepO* transcription was not affected by CSP in any carbohydrate tested. Therefore CSP does not stimulate *comX* by modulating *pepO* transcription. This finding is consistent with a model where CSP exerts a greater effect on *comR* while *pepO* acts through a separate mechanism to suppress competence activation.

As a test for possible effects of other genes proximal to the *tre* operon and *pepO*, we tested whether the transcription of *smu.2035*, which is located 143 bp downstream of *pepO* and transcribed in the opposite direction, was affected by CSP or carbohydrate. Fig. 4D shows that *smu.2035* transcripts decreased slightly in the Δ*pepO* background relative to the wild type. This may be a consequence of insertion of the antibiotic resistance cassette in the *pepO* region (*Methods)*. However, *smu.2035* showed no particular response to CSP or carbohydrate in the wild type.

### Deletion of *pepO* allows population-wide expression of *comX*

Figs. 3 and 4 show that deletion of *pepO* increased c*omS* transcription and transformability. However, *pepO* transcription is not modulated by CSP or carbohydrate. The simplest interpretation of these data is that the endopeptidase PepO acts constitutively to limit the activation of the ComRS autofeedback loop. We tested this model by measuring the proportion of cells activating a P*comX-gfp* reporter in the *pepO* deletion and in the wild-type background, in three different carbohydrates. Only a fraction (Fig. 5A, B) of wild-type cells became P*comX*-active at saturating concentrations of CSP, where this fraction was greater in trehalose or maltose (47 ± 3 % and 38 ± 5 % responding, respectively) than in glucose (22 ± 9 % of cells responding). The deletion of *pepO* substantially enhanced (Fig. 5C) the level of activation observed at saturating concentrations of CSP in all three carbohydrates: in trehalose and maltose, the proportion of *comX*-active cells approaches 100%, while in glucose it exceeds 90% (Fig. 5D). Therefore the bimodal behavior is largely removed by the deletion of *pepO* and the competence pathway now responds unimodally to CSP. Within each strain, the binding parameter *K* (Hill function with *n* = 1, Supplemental Table S1) was roughly the same for all sugars (*K* = 11-14 nM), indicating that the sugar does not determine the CSP sensitivity threshold. However the *K* for the Δ*pepO* strain was 2-4 nM, which is roughly 2-3-fold lower than in UA159. Therefore, deletion of *pepO* increased overall sensitivity to CSP in all three carbohydrates, and eliminated the bimodal character of *comX*, allowing population-wide activation of *comX*. Complementation of the *pepO* deletion (Fig. 5E) reduced the *comX*-active proportion to the level of 30-40% similar to the wild type and with a similar *K* = 11 ± 3 nM, reverting behavior to bimodal (Fig. 5F). These data show that PepO is a major limiting factor in the activity of the ComRS feedback loop, such that *pepO* deletion allows population-wide *comX* expression in the presence of CSP.

**Fig. 5:**
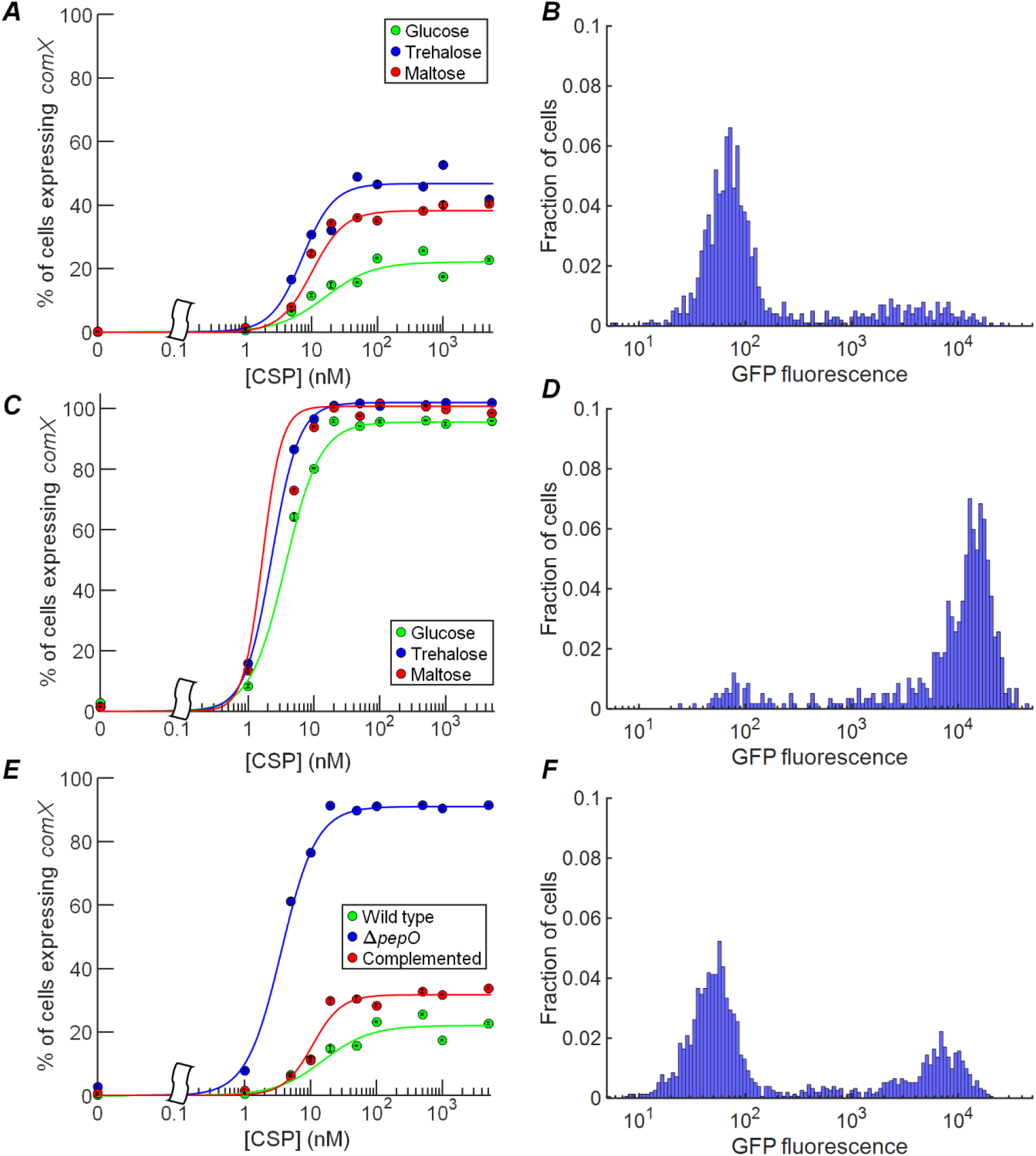
*pepO* deletion results in carbohydrate-dependent unimodal response to CSP. Percentage of P*comX-gfp* cells responding to CSP in TV medium supplemented with different sugars. Percentages are determined by comparing the area under the high and low-fluorescence peaks in the population distribution (see *Methods*). (A) Response of P*comX-gfp/*UA159 in glucose (green), trehalose (blue) or maltose (red). (B) Histogram of GFP fluorescence for individual cells provided with 5 μM CSP in glucose, demonstrating bimodal response. (C) Response of P*comX-gfp* Δ*pepO* strain in different carbohydrates, using same color code as in (A). (D) Single-cell histogram for P*comX-gfp* Δ*pepO* strain in glucose with 5 μM CSP, showing near unimodal (>90%) activation by CSP in glucose. (E) Comparison of the above P*comX-gfp/*UA159 (green) data and P*comX-gfp* Δ*pepO* (blue) data to a P*comX-gfp* Δ*pepO pepO*^+^ (complemented) strain (red), all grown in glucose. (F) Histogram of single-cell fluorescence of *pepO* complemented strain, grown in glucose with 5 μM CSP, showing restoration of the bimodal distribution seen in (B).

### PepO is responsible for growth medium-dependent bimodal response to CSP

We have proposed a mechanism where the growth medium dependence of the *comX* response to CSP is due to intracellular XIP/ComS and small nutrient peptides from the media competing for degradation by an intracellular peptidase (Son, M. *et al.*, 2012; Hagen; Son, 2017): In defined media that lack the peptides, the peptidase is available to degrade intracellular XIP/ComS, shutting down ComRS feedback and preventing *comX* activation. In complex media, the small peptides slow the degradation of XIP/ComS sufficiently to allow the ComRS feedback loop to autoactivate in some cells, leading to a population-bimodal *comX* response. If PepO plays the role of this hypothesized peptidase, then we would expect a *pepO* deletion strain to show an enhanced *comX* response to CSP in both complex and defined growth media. We therefore provided CSP to the Δ*pepO* strain in the defined medium FMC, supplemented with either glucose or trehalose. CSP normally elicits no response from *comX* in FMC medium (Son, M. *et al.*, 2012). However, Fig. 6 shows that even in the absence of CSP the Δ*pepO* strain was bimodally activated in both trehalose-supplemented FMC and in FMC formulated with glucose as the sole carbohydrate. Further, as the CSP concentration was increased to 100-500 nM (in glucose, Fig. 6A) or to 5-10 nM (in trehalose, Fig. 6B), the *comX* response became unimodal (population-wide). Therefore deletion of *pepO* eliminated both the bimodality of the *comX* response to CSP and the requirement for complex growth medium.

**Fig. 6:**
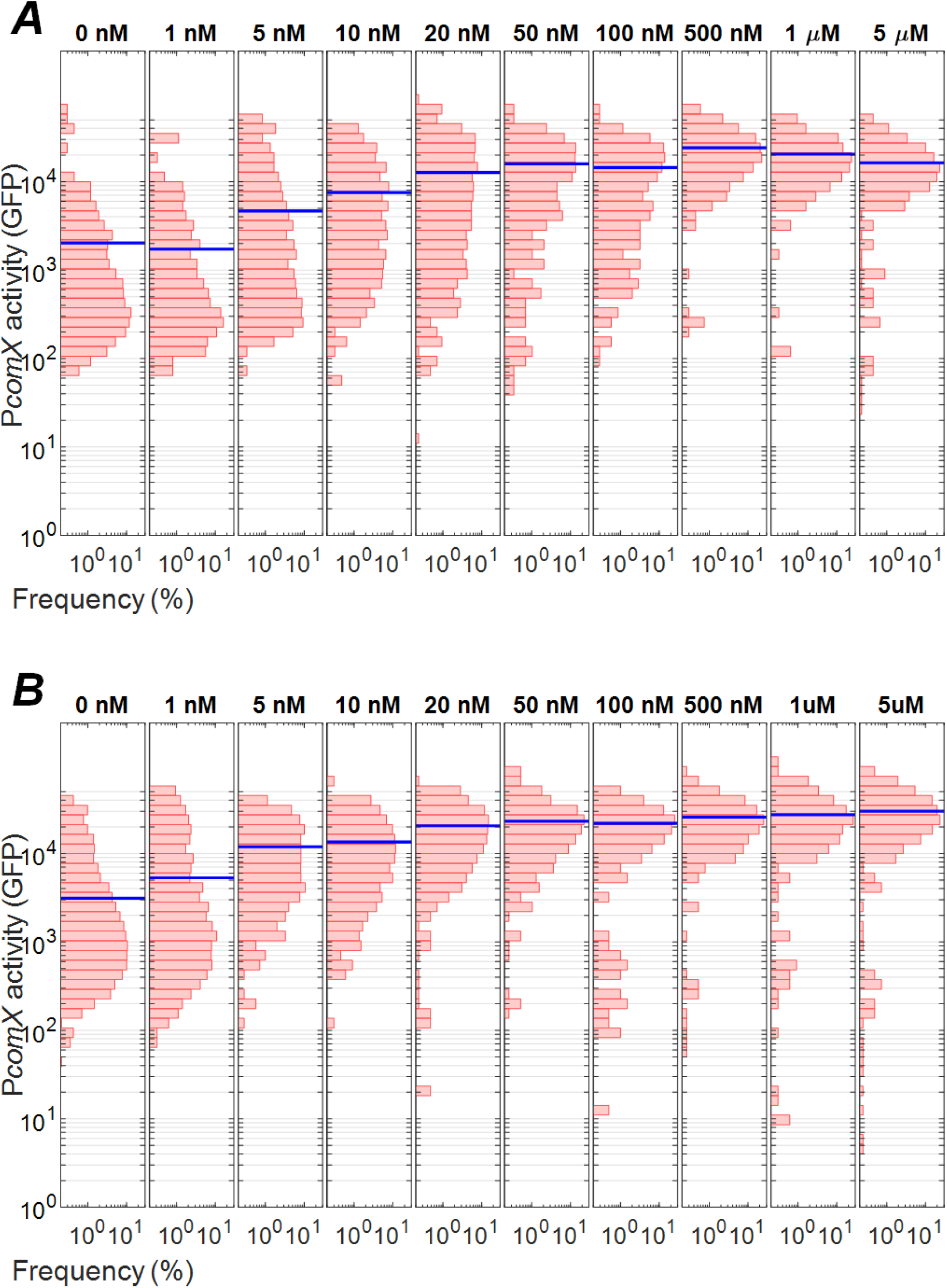
CSP-mediated activation of *comX* in a defined medium. P*comX-gfp* Δ*pepO* fluorescence response to indicated concentrations of added CSP in the defined medium FMC as measured by fluorescence microscopy in (A) glucose and (B) trehalose.

### PepO degrades the ComS-derived signal XIP *in vitro*

We have previously shown (Underhill *et al.*, 2018) that the bimodal response of ComRS and *comX* to CSP arises within an intracellular feedback loop in which endogenously produced ComS interacts with ComR to drive *comS* and *comX* transcription. The above data support the interpretation (Son, M. *et al.*, 2012) that the growth medium dependence of CSP response is due to a peptidase, evidently PepO, which suppresses autoactivation of the ComRS system by degrading the endogenous ComS feedback signal. In order to confirm that PepO can break down ComS/XIP and prevent its interaction with ComR to drive ComRS transcriptional feedback, we tested whether recombinant PepO (rPepO) from *S. mutans* affected the ability of synthetic XIP to form a DNA-binding complex with ComR *in vitro*. Fig. 7A shows a fluorescence polarization (FP) assay in which purified ComR binds a fluorescently labeled DNA probe containing the *comX* promoter region in the presence of 5 μM synthetic XIP that had been incubated with 500 nM rPepO for different lengths of time. A loss of polarization was observed when XIP was incubated for more than about 20 minutes, indicating a loss of XIP-induced binding of ComR to the DNA probe. A greater loss of polarization occurred at higher rPepO concentrations. Fig. 7B shows that 2 h incubation of rPepO with XIP had little effect on the FP signal if rPepO was present at concentrations below about 30 nM, but polarization declined substantially for [rPepO] greater than about 100 nM. Fig. 7C shows the FP signal for XIP that was incubated for 5 h at the same rPepO concentrations as Fig. 7B.

**Fig. 7:**
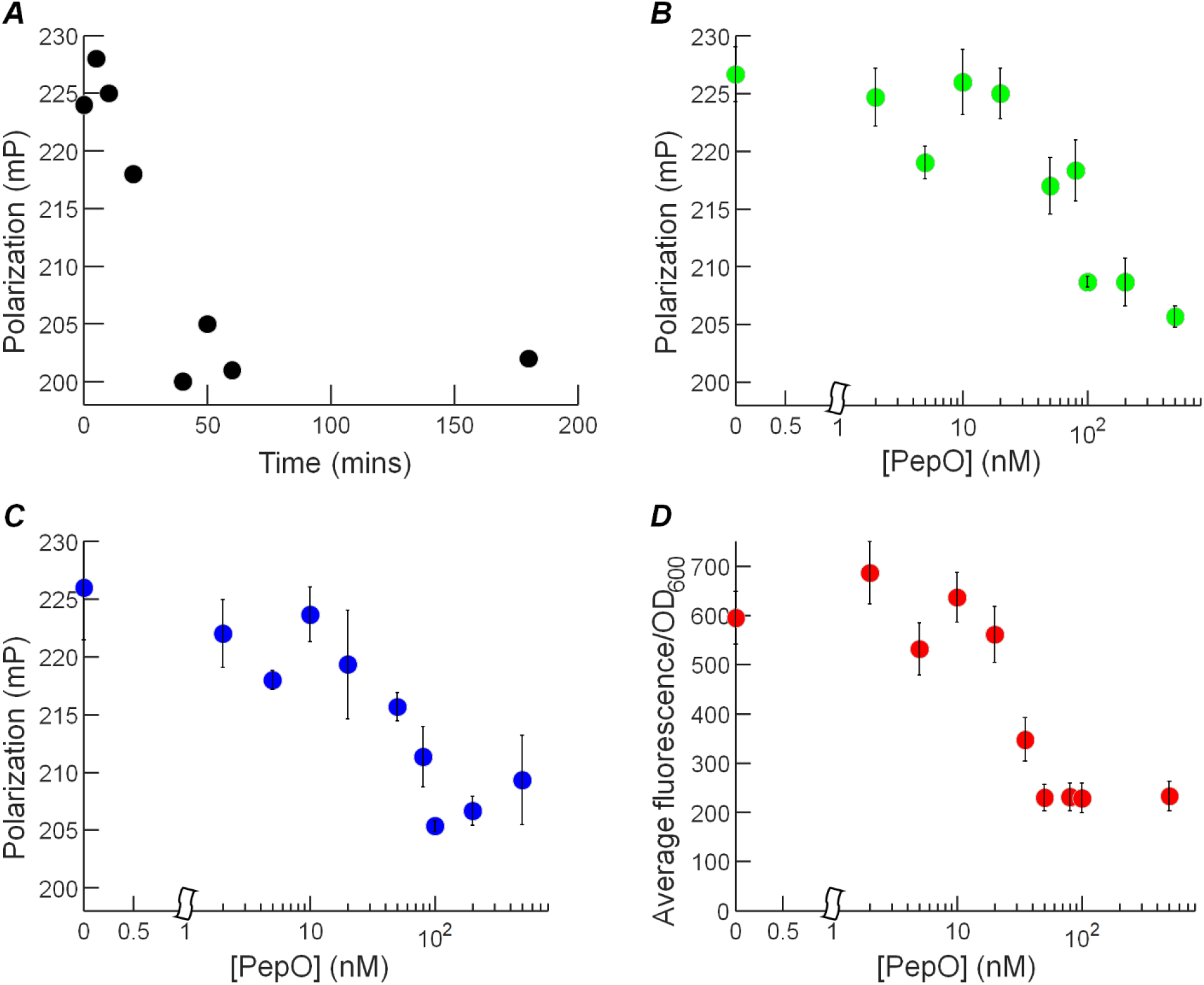
PepO degrades XIP *in vitro*. Effect of rPepO protein on synthetic XIP. (A) Effect of XIP incubation time with rPepO on a fluorescence polarization assay for XIP/ComR binding to the *comX* promoter. 500 nM rPepO was added to 5 μM synthetic XIP, and fluorescence polarization measurements were then taken after the indicated incubation times, using a fluorescently-labeled DNA aptamer and 1 μM purified recombinant ComR. (B), (C) Different concentrations of rPepO added to 5 μM synthetic XIP and fluorescence polarization measurements similar to (A) taken at 2 and 5 hours of incubation respectively. (D) Fluorescence response of P*comX-gfp* Δ*comS* cells to addition of rPepO-treated XIP added to culture in 1:4 dilution after 5 hours of rPepO treatment.

As an additional test of rPepO degradation of XIP, we tested whether treatment with rPepO affected the ability of synthetic XIP to activate *comX* in a reporting strain of *S. mutans*. Fig. 7D shows fluorescence of a bulk culture of P*comX-gfp* Δ*comS* cells (incapable of producing their own ComS or XIP) that were provided with 1 μM XIP that had been incubated with rPepO for 5 hours. The reporter fluorescence of the cells shows a decline very similar to Fig. 7C as the rPepO concentration is increased, confirming that rPepO degraded the ability of the XIP to activate *comX*.

## Discussion

Genetic competence in *S. mutans* is influenced by a diverse set of environmental factors, including peptide content of the medium, pH, oxidative stress and heat (Senadheera, M. Dilani *et al.*, 2005; Ahn *et al.*, 2006; Tremblay *et al.*, 2009; Okinaga *et al.*, 2010; Son, M. *et al.*, 2012; Guo *et al.*, 2014; De Furio *et al.*, 2017). For *S. mutans*, the peptide content of the growth medium determines the proportion of cells that activate *comX* in response to CSP. The response ranges from zero (in defined medium, lacking small peptides) to partial (bimodal, in complex media). Only direct addition of the inducing peptide XIP, which when taken up by the Opp oligopeptide permease in defined media interacts directly with ComR to induce *comX*, elicits a population-wide (unimodal) *comX* response in wild-type *S. mutans*. Moye *et al.* (Moye *et al.*, 2016) showed that carbon source is an additional modulator of the proportion of cells responding to CSP by activating *comX*. In particular, growth in the presence of the disaccharide trehalose not only increases the proportion of *S. mutans* expressing *comX*, but also increases transformation efficiency. The trehalose catabolic operon is directly upstream of, and transcribed in the same direction as, the endopeptidase *pepO*. In investigating the link between carbohydrate source and bimodal activation of *comX*, we initially hypothesized that the proximity of *pepO* and the *treAB* operon may allow induction of *treAB* to modify levels of the peptidase. This could lead to higher intracellular ComS, encouraging autoactivation of the bistable ComRS feedback circuit that regulates *comX* (Fig. 1) (Son, M. *et al.*, 2012). Although our findings confirm the trehalose effect originally observed by Moye et al. (Moye *et al.*, 2016) and demonstrate for the first time that PepO plays a key role in modulating bimodal competence response, the body of our data shows that carbohydrate source and PepO influence the regulatory pathway through largely independent mechanisms.

It was necessary to determine whether the trehalose effect originates with the bacteriocin genes because the trehalose operon has been linked to bacteriocin expression (Baker *et al.*, 2018). Our data rule out a role for *cipB* or upstream elements such as *comCDE* in the effect of trehalose and other carbohydrates tested, because CSP activation of *cipB* was unaffected by carbohydrate source. The CSP concentration required to saturate the *cipB* response, and the maximal level of *cipB* expression, was similar in glucose and trehalose.

Figure 8A shows further evidence that carbohydrate affects the competence pathway downstream of *cipB*. Expression of *cipB* reaches maximum at a lower CSP concentration (near 5 nM CSP) than does expression of the *PcomX-gfp* reporter (near 100 nM CSP). Therefore, even though *cipB* is required to elicit the *comX* response in glucose (Perry *et al.*, 2009), trehalose and maltose (Supplemental Figure S2), full activation of *cipB* is insufficient to induce maximum *comX* response. Evidently, transmission of the competence signal beyond *cipB* involves an additional mechanism that requires a higher threshold of CSP, such as activation of an additional gene that is subject to *comDE* regulation. The carbohydrate source does not affect the threshold CSP concentration (constant *K* in Table S1) needed to saturate either *cipB* or *comX* expression. Therefore, the carbohydrate effect on *comX* appears to arise in an additional mechanism unlinked to *cipB*, whereby CSP leads to upregulation of *comR* transcription, and where glucose appears to inhibit or interfere with this mechanism.

**Fig. 8:**
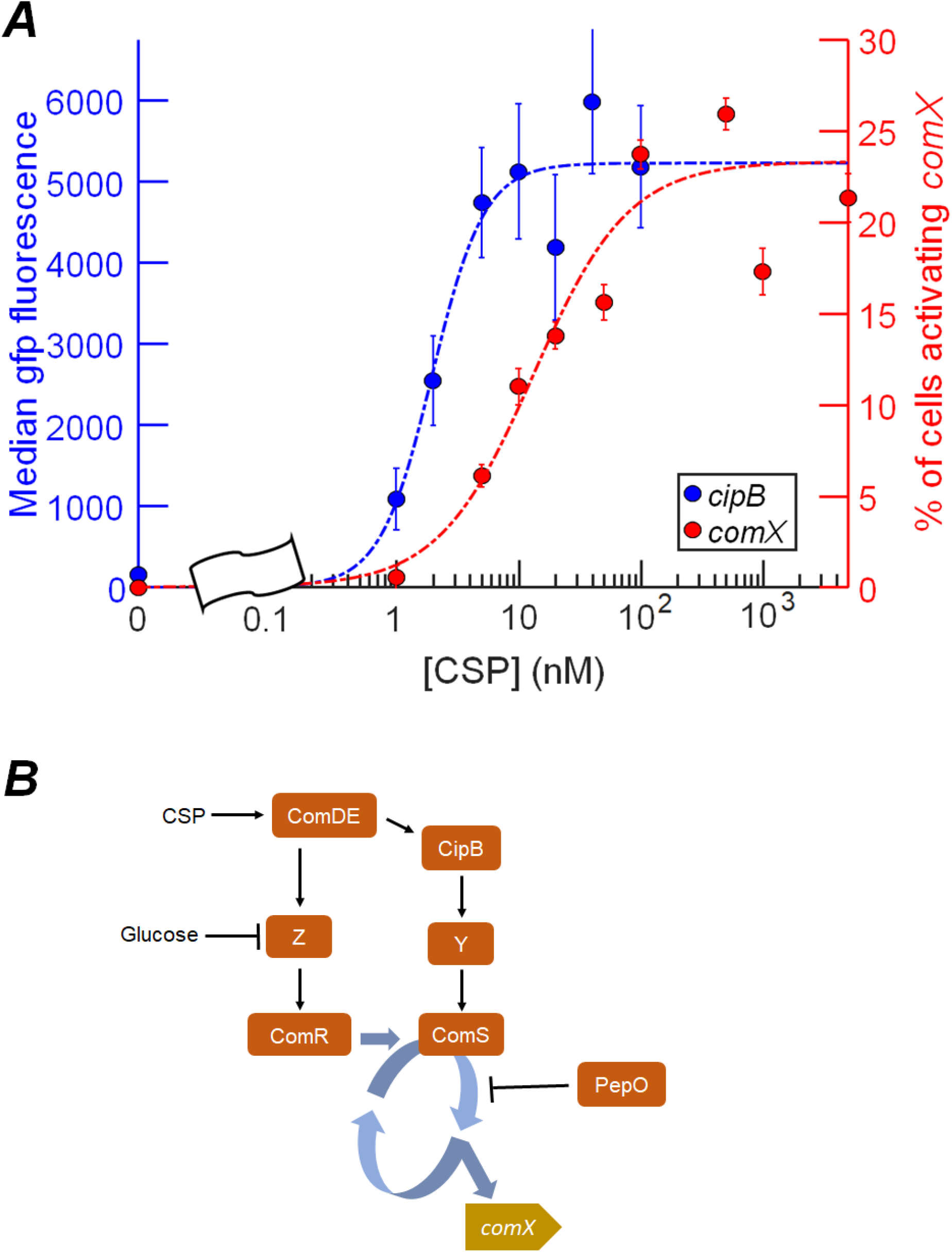
integration of carbohydrate effects into existing competence regulation models. (A) P*cipB-gfp* and P*comX-gfp* expression data from Figs. 2 and 5 (glucose, wild type) on a double y-axis plot showing the difference in [CSP] threshold between *comX* and *cipB*. (B) Model for the influence of carbohydrate on the *comX*-activating fraction of the population, in response to CSP. An unknown mechanism Z increases *comR* transcription under stimulation by CSP, but is repressed by glucose, such that *comR* is only significantly upregulated in the other carbohydrates tested. A parallel pathway Y is also postulated, through which phosphorylated ComE is able to stimulate *comS* via expression of *cipB*. The response of *comS* to CSP stimulation requires the autofeedback amplification of fluctuations in [ComS], which is repressed by PepO-mediated degradation, limiting the proportion of the population that activates *comX*.

Our data indicate that PepO affects the competence pathway in a different way than carbohydrate, most likely by degrading basally produced ComS and XIP. This action inhibits the transcriptional feedback that amplifies fluctuations in ComS levels into full activation of ComRS and induction of *comX.* If the effect of *comCDE* stimulation is to increase *comS* or *comR* transcription, then it will help to overcome the suppressive effect of PepO and permit, in at least some cells, self-activation of the feedback loop and flipping of the ComRS switch to its ON state. Consistent with this model, our transcriptional data suggest that in the tested carbohydrates (but not in glucose), CSP leads to some upregulation of *comR*. Although *comR* transcripts increase only be a modest 2- to 3-fold, bistable autofeedback systems amplify small fluctuations (Dubnau; Losick, 2006) and a small rise in ComR levels could very plausibly flip some cells from the *comX*-OFF to the *comX*-ON state.

PepO is very strongly implicated in the role of feedback inhibitor because its deletion drastically enhanced transformability and triggered population-wide expression of *comX*. A robust, unimodal *comX* response to CSP, which has not otherwise been observed in *S. mutans*, is then observed even in defined media, where CSP normally elicits no response from *comX* whatsoever (Son, M. *et al.*, 2012) regardless of carbohydrate source (Ricomini *et al.*, 2019). Degradation of ComS/XIP by PepO is also consistent with observations such as the degradation of the RGG2 and three other small peptides by PepO in *Streptococcus pyogenes* (Wilkening *et al.*, 2016). As PepO amino acid sequences are ~90% similar across *Streptococcus* spp. (Nguyen *et al.*, 2009), the relatively non-specific degradation of small signal peptides seen in (Alves *et al.*, 2017) is potentially a conserved property of PepO peptidase. Our *in vitro* data support this interpretation by showing that pre-treatment of XIP or ComS with PepO inhibits ComR binding to its cognate target, apparently by degrading exogenously added XIP. We also showed that the effect of PepO on ComR/XIP-dependent activation of *comX* can be reproduced *in vivo* where treatment of XIP with rPepO prior to addition to ComS-deficient cells in defined medium eliminates the ability of XIP to activate *comX* in cells.

It appears unlikely that PepO plays a direct role in the carbohydrate effect, as *pepO* expression was not modulated by CSP or carbohydrate source. PepO is not controlled through the ComDE circuit and instead acts independently to suppress autoactivation of ComRS. The model presented in Fig. 8B therefore proposes that PepO acts constitutively to degrade endogenously produced ComS and nutritional peptides from the medium. The model also includes a parallel, CSP-dependent pathway – denoted Z – that stimulates *comR* transcription in a CSP-dependent manner, but is sensitive to carbohydrate source. The efficiency with which CSP is capable of activating *comR* is thus controlled by the sensitivity of Z to CSP and by carbohydrate source, as our RT-qPCR data indicate in Fig. 4D.

We note that prior transcriptional studies have disagreed on whether or not CSP increases ComR levels (Lemme *et al.*, 2011; Reck *et al.*, 2015; Moye *et al.*, 2016). Our data and model resolve the apparent disagreement, inasmuch as *comR* upregulation was not observed when CSP was provided in glucose-supplemented chemically defined medium (Reck *et al.*, 2015), but a 1.4- to 1.8-fold upregulation was detected in THB-Y medium (Lemme *et al.*, 2011), which contains other carbohydrates, in addition to glucose, that are presumably able to trigger the carbohydrate-sensitive pathway.

Our data still leave unanswered the question of how CSP stimulates ComRS when glucose is the sole carbohydrate source and ComR levels are unaffected by CSP. Here, it may be relevant to observe that although the *pepO* mutant growing in the presence of CSP and glucose activates *comX* far more robustly than in a wild-type genetic background (Fig. 5C and Fig. 6A), this response still requires a small (5-10 nM) concentration of CSP. Even in the absence of PepO, the ComRS feedback system is still weakly repressed, although only modest amounts of CSP are needed to overcome this repression. Similarly, *comX* does not respond to CSP when a *cipB* deletion strain grows in the disaccharides tested (Supplemental Fig. S1). All of these results indicate that *cipB* controls a pathway (denoted Y in Fig. 8B) that operates in all growth media to limit ComRS activation, independently of Z. It is conceivable for example that an additional peptidase (other than PepO) weakly degrades endogenously produced ComS/XIP, and that the *cipB* pathway acts to downregulate this peptidase. Such a model is depicted in Fig. 8B, where CSP has two parallel effects on ComRS: it downregulates the second protease (pathway Y) while also upregulating *comR* (pathway Z). This model predicts that in a *pepO* deletion strain in glucose-containing media (Z not active), the CSP threshold for the *comX* response will be similar to that for *cipB* activation – as occurs in our data (Fig. 2 and Fig. 5). However, for UA159 growing in other carbohydrates, the CSP level that is needed to induce such a pathway will be difficult to predict, owing to the two parallel routes combined with the internal positive feedback, which is present both in ComRS (Son, M. *et al.*, 2012; Underhill *et al.*, 2018) and in regulation of *comDE* by ComX (Son, Minjun *et al.*, 2015; Reck *et al.*, 2015).

Finally, our results allow some speculation about how the VicRKX sensory system, which is hypothesized to respond to oxidative stress (De Furio *et al.*, 2017), links to the competence pathway. A binding site for the VicR response regulator has been identified in the *pepO* promoter region (Alves *et al.*, 2017), where VicR has been demonstrated to act as a transcriptional repressor (Senadheera, D. B. *et al.*, 2012). Thus, the *vicK* deletion results in higher *pepO* expression (Alves *et al.*, 2017), which should give rise (under our model) to reduced transformability in these mutants. A prior study showed in fact that deletion of the *vicK* kinase reduces transformability, despite increasing the production of bacteriocins and *comCDE* mRNA (Senadheera, M. Dilani *et al.*, 2005). A similar pattern is visible in data showing that deleting *clpP* raises *pepO* expression while concurrently lowering *comR* and *comX* expression (Kajfasz *et al.*, 2011).

## Acknowledgments

The authors acknowledge the NIDCR for funding through 1R01DE023339, 1R01DE13239 and 1R01DE12236. We also extend our gratitude to Dr. Livia Alves of the University of Florida College of Dentistry for the kind gift of rPepO protein and Dr. Jessica Kajfasz at the University of Florida College of Dentistry for the Δ*pepO pepO*^+^ complemented strain.

## Materials and Methods

### Strains and growth conditions

*S. mutans* wild-type strain UA159 and mutant strains from glycerol freezer stocks were grown in BBL BHI (Becton, Dickinson and co.) at 37°C in 5% CO_2_ overnight. *E. coli* were grown from glycerol freezer stocks in LB at 37°C shaking overnight. Antibiotics were used at the following concentrations where resistance is indicated in Table 1: erythromycin (10 μg ml^−1^), kanamycin (1 mg ml^−1^), spectinomycin (1 mg ml^−1^), ampicillin (10 μg ml^−1^). For all experiments, strains were washed twice by centrifugation, removal of supernatant fluids and re-suspension in phosphate buffered saline (PBS), pH 7.2. Cells were then diluted 20-fold into fresh medium and allowed to grow in the same incubator conditions until OD_600_ reached 0.1. Synthetic CSP-18 (sequence SGSLSTFFRLFNRSFTQA) was purified to 98% purity and provided by NeoBioSci (Cambridge, MA, USA).

**Table 1:**
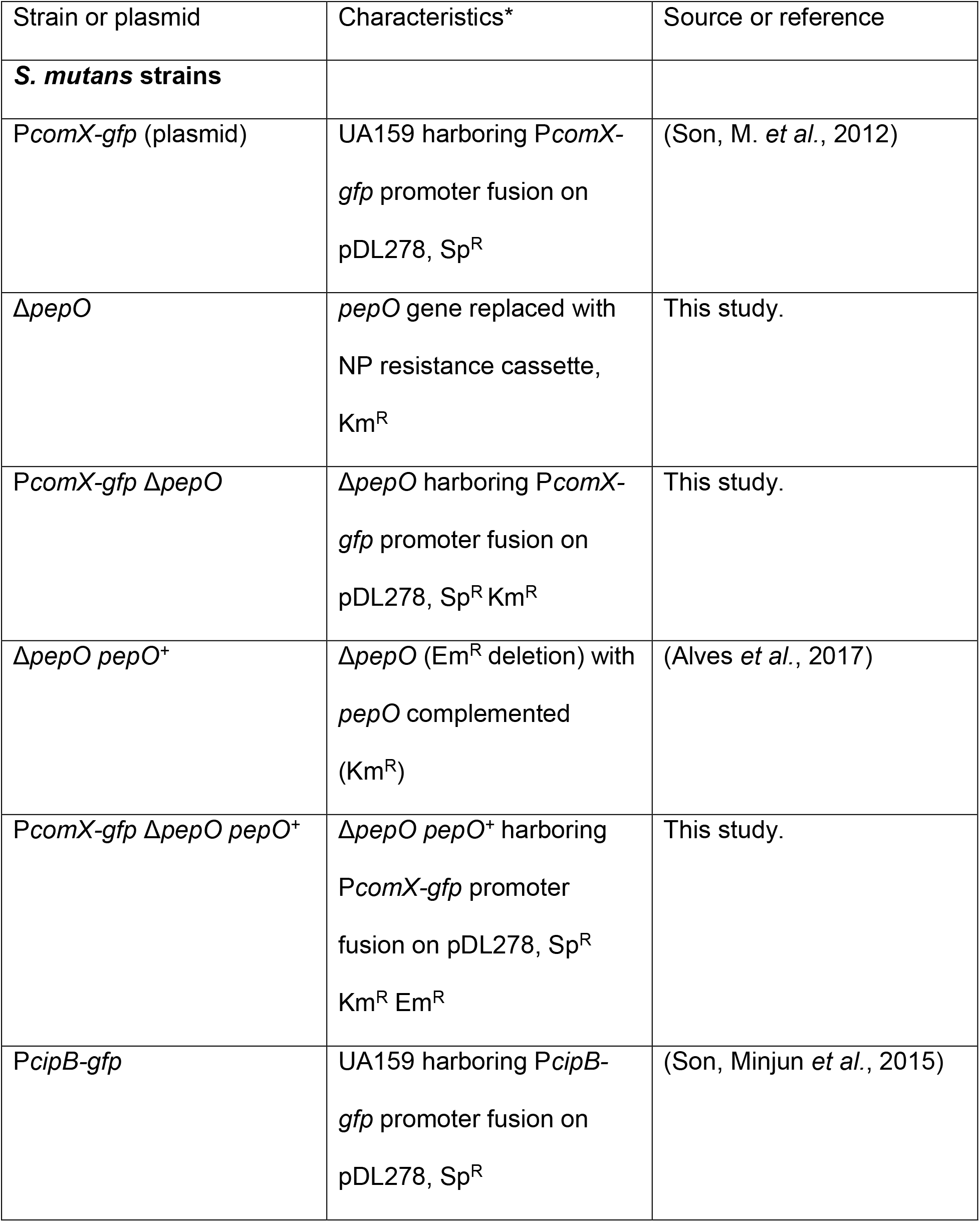
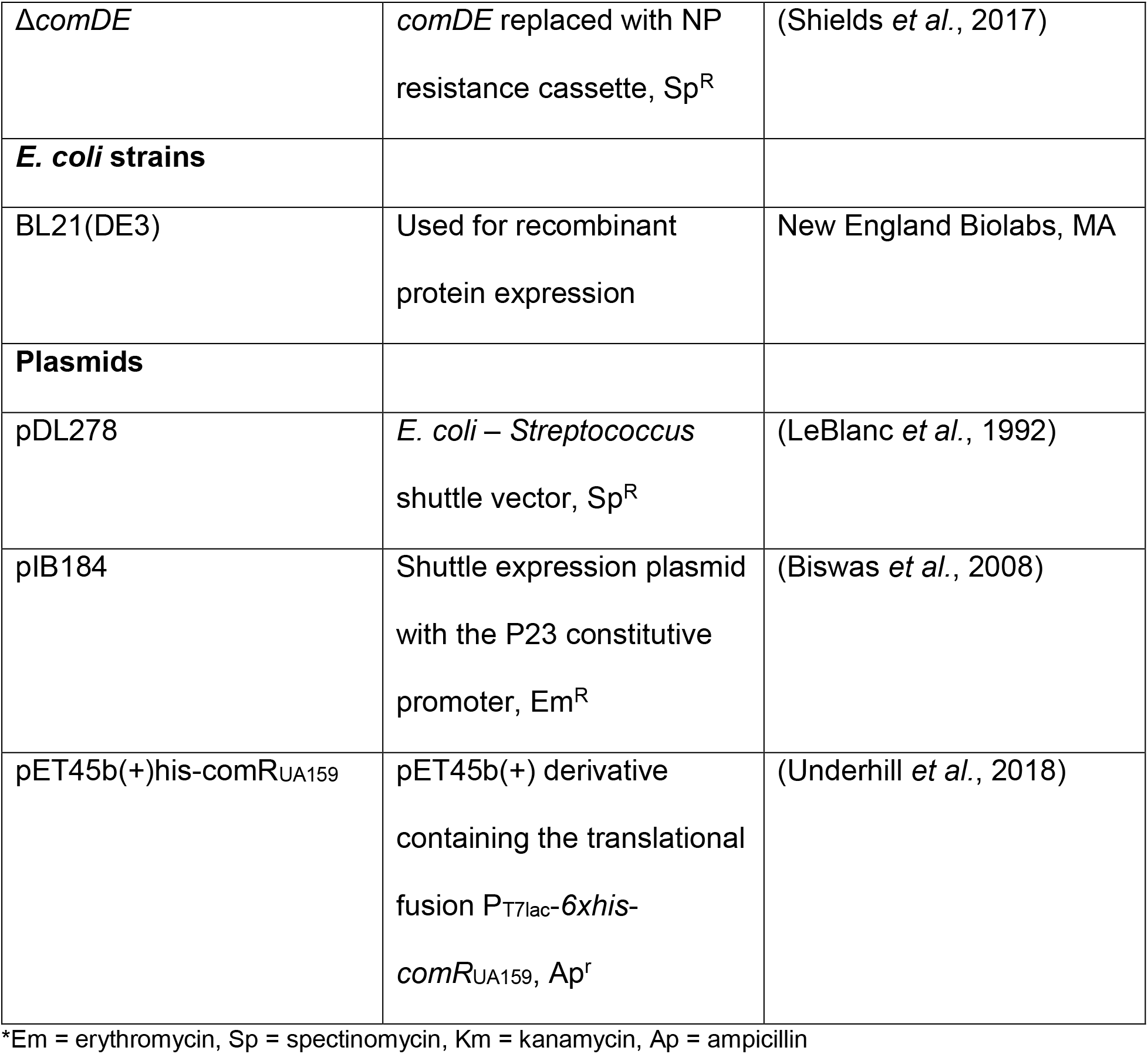
strains and plasmids used. List of strains and plasmids used in this study, their relevant characteristics, and the source or reference for the material.

### Construction of *pepO* deletion mutant

The *pepO* gene was replaced by a non-polar kanamycin cassette in *S. mutans* strain UA159 by homologous recombination. PCR primers (Table 2) with ends containing a *Bam*HI recognition site were used to amplify the flanking regions of the gene. Ends were digested with *Bam*HI to ligate the flanking region product to the kanamycin cassette. The resulting linear DNA was transformed into UA159 in which competence was induced by XIP in the defined medium FMC (Terleckyj *et al.*, 1975). The transformants were confirmed by PCR and Sanger sequencing to ensure that the *pepO* gene was deleted and the sequences flanking *pepO* that were used for the recombination event were intact. Previously described fluorescent protein reporter fusion constructs (Son, M. *et al.*, 2012) were transformed as needed to generate strains for use in experiments.

**Table 2:**
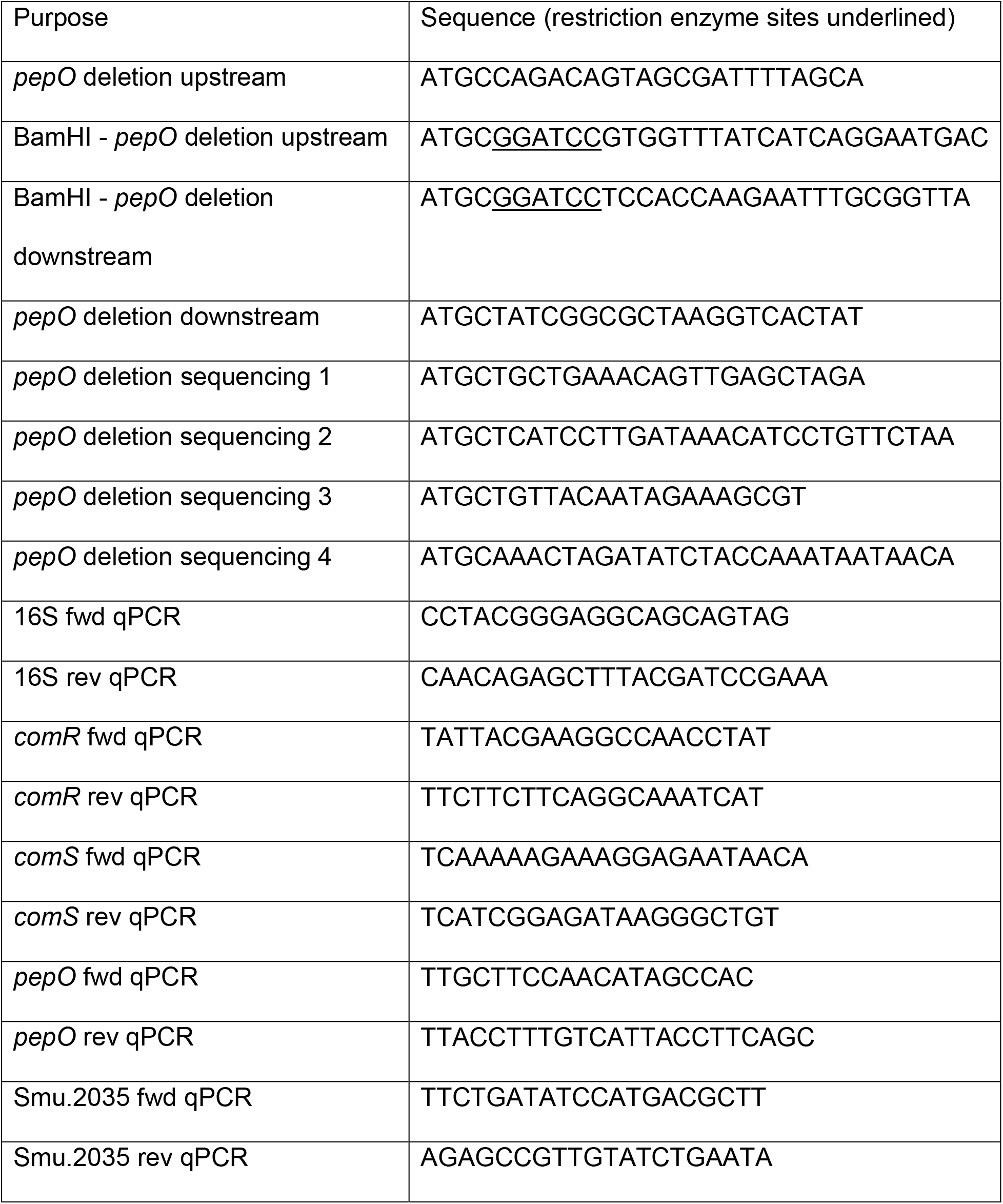
oligonucleotides used. Primers used for this study, including qPCR primers. Restriction enzyme sites are underlined.

### Development of buffered TV medium

The pH of tryptone-vitamin (TV) medium was found to be approximately 6.7. Due to the influence of pH on the *com* regulon (Guo *et al.*, 2014; Son, M. *et al.*, 2015), a medium buffered at a pH close to 7 was desired. A phosphate buffer (170 mM KH_2_PO_4_, 720 mM K_2_HPO_4_) was diluted into TV medium 100-, 80-, 50- or 10-fold and cells from overnight cultures were diluted 20-fold into the differently buffered media. The resulting cell suspensions plus an unbuffered control were put into a well plate (Falcon 24 well plate, Corning inc.) and growth was monitored by measuring the OD_600_ every 5 minutes in a Biotek Synergy 2 plate reader (Biotek Instruments, inc.). It was found that the 80-fold dilution was the highest concentration of buffer that did not inhibit growth of *S. mutans* (Supplemental Figure S3) so medium buffered in this was used for all carbohydrate experiments involving TV. The initial pH of the buffered TV was 7.2, which is known to be permissive for competence signaling by CSP (Guo *et al.*, 2014; Son, M. *et al.*, 2015).

### Single cell experiments

For single-cell experiments involving planktonic growth in different carbohydrates in the presence of CSP, a chromosomally integrated P*comX-gfp* reporter strain was used. Cells were diluted 20-fold from overnight cultures into buffered TV supplemented with the desired carbohydrate(s) at final concentrations of 20 mM for monosaccharides and 10 mM for disaccharides. When the cells reached an OD_600_ of 0.1, 1 μM CSP-18 was added and the cultures were incubated for 2 h. At this point, cultures were gently sonicated using a Fisher Scientific FB120 sonic dismembrator probe to break up long chains and pipetted onto a glass coverslip. Phase contrast and fluorescence imaging were performed at 60x magnification on a Nikon TE2000U phase contrast microscope followed by image analysis, as described previously (Kwak *et al.*, 2012). The percentage of cells deemed to be expressing *comX* was determined using a two-distribution fit to the bimodal data (see below).

### Concurrent monitoring of growth and fluorescence

Well plate experiments were carried out in a Falcon 96 well plate with clear bottom and black side (Corning, Inc.). 200 μL of culture was pipetted into each well and covered with mineral oil to prevent evaporation and oxygen diffusion. Fluorescence and OD_600_ were read in a Biotek Synergy 2 plate reader (Biotek Instruments).

### Transformation efficiency assay

Cells were prepared for each transformation efficiency assay from overnight cultures as for the CSP experiments described above. DNA used for transformation was plasmid pIB184, carrying an erythromycin (Em) resistance marker, added at a concentration of 600 ng ml^−1^. At OD_600_ of 0.1, DNA was added to the cells and tubes were mixed by inversion; at this point CSP was added where used. After 4 hours of exposure to DNA, cells without CSP (or *comDE* mutants in all cases, as these do not respond to CSP (Perry *et al.*, 2009)) were diluted and plated on BHI supplemented with 10 μg ml^−1^ erythromycin. A no-DNA control from each sugar was plated on BHI-erythromycin in order to verify that no spontaneously resistant variants arose during the incubation. Total viable cell counts were obtained by plating the diluted cultures onto BHI agar with no antibiotics. Efficiency was calculated by dividing the number of transformants per microliter plated by the total viable cell count per microliter plated, correcting for dilution and concentration factors appropriately.

### Calculating the *comX*-active proportion of cells

The double peaked (bimodal) distribution of P*comX* activity under CSP stimulation was found empirically to be well characterized as a combination of two well-separated gamma distribution functions (Friedman *et al.*, 2006; Taniguchi *et al.*, 2010) (as in Figure 1), corresponding to *comX*-ON and OFF cells respectively. For cells responding bimodally to CSP, the histogram *P(F)* of P*comX* fluorescence *F* can then be fit to obtain a parameter λ (0 ≤ λ ≤ 1) equal to the proportion of cells in which *comX* is active:

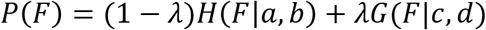

Here *H* is the gamma distribution with shape parameters *a* and *b*, describing the OFF cells, and *G* is the gamma distribution with parameters *c* and *d*, describing the ON cells. In experiments when P*comX* activity was nearly unimodal (single peaked), this fit did not determine all parameters robustly, and so the *comX*-active proportion λ was found by a cutoff method, based on counting the proportion of cells whose fluorescence *F* exceeds a cutoff value *F*_*c*_. We chose the cutoff *F*_*c*_ by identifying the value that, for a comparable dataset where P*comX* activity is strongly bimodal, minimizes the probability that a simple cutoff wrongly assigns a cell to either the ON or OFF distribution. That is, the fluorescence cutoff *F*_*c*_ was chosen to minimize

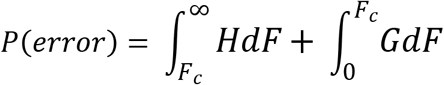

Hill functions were fit to data exhibiting saturating behavior using a Hill function model with *n* set to 1 and hence two free parameters, the saturating height and *K*. Fits were performed by the chi squared nonlinear fitting method and error in parameters estimated by a 1000 iteration bootstrap for each data set.

### *In vitro* degradation of XIP by PepO

Purified recombinant PepO (rPepO) was a kind gift of Livia Alves and Dr. Jacqueline Abranches of the University of Florida College of Dentistry. Purified protein was frozen in PBS pH 7.2 with 10% glycerol. The concentration of rPepO in solution was estimated from its absorbance at 280 nm. XIP was treated with rPepO by diluting XIP from 10 mM phosphate buffer at pH 7.0 (5.8 mM K_2_HPO_4_, 4.2 mM KH_2_PO_4_) with indicated concentrations of rPepO. The mixtures were then incubated at 37°C for 5 hours before addition to cells.

To detect how much XIP was left, the rPepO-XIP mixtures were added to P*comX-gfp ΔcomS* plasmid-based reporter cells at OD_600_ 0.1 in a ratio of 1:4 (such that the XIP mixture was 5-fold diluted into the cell culture) in a 96-well plate and OD_600_ and green fluorescence was recorded every 5 min. The GFP fluorescence was normalized to OD_600_ and averaged over the same time period for each sample corresponding to the time between onset and decline of the fluorescence peak. The data were plotted using the standard deviation of F/OD_600_ values over this time as error bars.

### Fluorescence polarization using rPepO-treated XIP and ComS

Native ComR was purified by previously reported methods (Underhill *et al.*, 2018). Briefly, *E. coli* BL21 (DE3) cells containing a plasmid harboring N-terminally tagged 6x His-ComR were lysed and the ComR purified by Ni-NTA affinity chromatography. The histidine tag was cleaved using EKMax enterokinase (Invitrogen), and the resulting ComR dialyzed into PBS pH 7.4. Concentration was estimated using the Pierce bicinchonic acid (BCA) assay (Thermo Scientific). The purified protein was used with the same Bodipy-FL-X labeled fluorescent DNA aptamer and binding assay buffer as described elsewhere (Underhill *et al.*, 2018) to assess the ability of rPepO-treated XIP to induce ComR binding of the P*comX* DNA region.

XIP (5 μM) was treated with the indicated concentrations of rPepO in phosphate buffer. After 2 and 5 hours, 40 μL of the XIP-rPepO solutions were pipetted into a well containing 160 μL of binding buffer, 1 nM fluorescent DNA and sufficient ComR to obtain 1 μM final concentration in the 200 μL volume. Fluorescence polarization was then measured by a Biotek Synergy 2 plate reader (Biotek Instruments inc.) in a 96-well black-bottomed, black-side assay plate in polarization mode using a 485 nm excitation filter and 528 nm emission filter. The same experiment was then performed using 5 μM XIP and 500 nM rPepO and taking polarization measurements at the indicated time points.

### Reverse transcriptase-quantitative PCR (RT-qPCR)

Cells for RT-qPCR were grown to OD_600_ = 0.1 in TV supplemented with the indicated sugar from 20-fold dilution from washed overnight cultures. At this point, CSP was added to samples and the cultures were incubated for 2 h before centrifugation and removal of the TV. One ml of TRIzol reagent (ThermoFisher) was then added, the pellet was resuspended, and cells were mechanically lysed in a bead beater in the presence of 100 μm glass beads. Extraction of RNA was then performed following the TRIzol phenol:chloroform method (Chomczynski, 1993) and the resulting RNA sample treated with the Turbo DNA-*free*™ kit from ThermoFisher.

RNA concentration was estimated using absorbance at 260 nm and 1 μg was reverse transcribed using iScript reverse transcription mix (Biorad) containing random primers. Resulting cDNA was diluted 50-fold in water and used as the basis for qPCR reactions. qPCR was performed using iTaq™ SYBR Green Supermix (Biorad) in a Biorad CFX Connect thermal cycler. Transcript counts for genes of interest were normalized to a count of 16S rRNA for the same volume of sample. Three biological replicates of each condition were grown and three independent technical replicates assayed for each of these. Resulting values of transcript count divided by 16S rRNA count were averaged to compute reported values. Standard deviations of mRNA counts in technical replicates were propagated forward to the calculated quotient and the computed error used to represent error bars.

### CSP activity in defined medium

To examine the influence of CSP on *comX* expression in a *pepO* mutant in defined medium, FMC, was supplemented with the indicated sugar (20 mM for glucose, 10 mM for trehalose) (Terleckyj *et al.*, 1975). The experiment was otherwise performed exactly as the single cell experiments in TV medium.

## References

Ahn, S.J., Lemos, J.A., and Burne, R.A. (2005) Role of HtrA in growth and competence of *Streptococcus mutans* UA159. J Bacteriol 187: 3028–3038.

Ahn, S.J., Wen, Z.T., and Burne, R.A. (2006) Multilevel control of competence development and stress tolerance in *Streptococcus mutans* UA159. Infect Immun 74: 1631–1642.

Alves, L.A., Harth-Chu, E., Palma, T.H., Stipp, R.N., Mariano, F.S., Höfling, J.F., et al. (2017) The two-component system VicRK regulates functions associated with *Streptococcus mutans* resistance to complement immunity. Mol oral Microbiol 32: 419–431.

Baker, J.L., Lindsay, E.L., Faustoferri, R.C., To, T.T., Hendrickson, E.L., He, X., et al. (2018) Characterization of the trehalose utilization operon in *Streptococcus mutans* reveals that the TreR transcriptional regulator is involved in stress response pathways and toxin production. J Bacteriol 200:e00057–18:.

Biswas, I., Jha, J.K., and Fromm, N. (2008) Shuttle expression plasmids for genetic studies in *Streptococcus mutans*. Microbiology 154: 2275–2282.

Chomczynski, P. (1993) A reagent for the single-step simultaneous isolation of RNA, DNA and proteins from cell and tissue samples. BioTechniques 15: 532.

De Furio, M., Ahn, S.J., Burne, R.A., and Hagen, S.J. (2017) Oxidative Stressors Modify the Response of *Streptococcus mutans* to Its Competence Signal Peptides. Appl Environ Microbiol 83: e01345–17.

Dubnau, D., and Losick, R. (2006) Bistability in bacteria. Mol Microbiol 61: 564–572.

Fontaine, L., Goffin, P., Dubout, H., Delplace, B., Baulard, A., Lecat-Guillet, N., et al. (2013) Mechanism of competence activation by the ComRS signalling system in streptococci. Mol Microbiol 87: 1113–1132.

Fontaine, L., Wahl, A., Fléchard, M., Mignolet, J., and Hols, P. (2015) Regulation of competence for natural transformation in streptococci 33: 343–360.

Friedman, N., Cai, L., and Xie, X.S. (2006) Linking stochastic dynamics to population distribution: an analytical framework of gene expression. Phys Rev Lett 97: 168302.

Guo, Q., Ahn, S.J., Kaspar, J., Zhou, X., and Burne, R.A. (2014) Growth phase and pH influence peptide signaling for competence development in *Streptococcus mutans*. J Bacteriol 196: 227–236.

Hagen, S.J., and Son, M. (2017) Origins of heterogeneity in *Streptococcus mutans* competence: interpreting an environment-sensitive signaling pathway. Phys Biol 14: 015001.

Hossain, M.S., and Biswas, I. (2012) An extracelluar protease, SepM, generates functional competence-stimulating peptide in *Streptococcus mutans* UA159. J Bacteriol 194: 5886–5896.

Kajfasz, J.K., Abranches, J., and Lemos, J.A. (2011) Transcriptome analysis reveals that ClpXP proteolysis controls key virulence properties of *Streptococcus mutans*. Microbiology 157: 2880–2890.

Kwak, I.H., Son, M., and Hagen, S.J. (2012) Analysis of gene expression levels in individual bacterial cells without image segmentation. Biochem Biophys Res Commun 421: 425–430.

LeBlanc, D.J., Lee, L.N., and Abu-Al-Jaibat, A. (1992) Molecular, genetic, and functional analysis of the basic replicon of pVA380-1, a plasmid of oral streptococcal origin. Plasmid 28: 130–145.

Lemme, A., Grobe, L., Reck, M., Tomasch, J., and Wagner-Dobler, I. (2011) Subpopulation-specific transcriptome analysis of competence-stimulating-peptide-induced *Streptococcus mutans*. J Bacteriol 193: 1863–1877.

Li, Y.H., Tang, N., Aspiras, M.B., Lau, P.C., Lee, J.H., Ellen, R.P., and Cvitkovitch, D.G. (2002) A quorum-sensing signaling system essential for genetic competence in *Streptococcus mutans* is involved in biofilm formation. J Bacteriol 184: 2699–2708.

Loesche, W.J. (1986) Role of *Streptococcus mutans* in human dental decay. Microbiol Rev 50: 353–380.

Mashburn-Warren, L., Morrison, D.A., and Federle, M.J. (2010) A novel double-tryptophan peptide pheromone controls competence in *Streptococcus* spp. via an Rgg regulator. Mol Microbiol 78: 589–606.

Moye, Z.D., Son, M., Rosa-Alberty, A.E., Zeng, L., Ahn, S.J., Hagen, S.J., and Burne, R.A. (2016) Effects of Carbohydrate Source on Genetic Competence in *Streptococcus mutans*. Appl Environ Microbiol 82: 4821–4834.

Nguyen, T., Zhang, Z., Huang, I., Wu, C., Merritt, J., Shi, W., and Qi, F. (2009) Genes involved in the repression of mutacin I production in *Streptococcus mutans*. Microbiology 155: 551–556.

Okinaga, T., Niu, G., Xie, Z., Qi, F., and Merritt, J. (2010) The hdrRM operon of *Streptococcus mutans* encodes a novel regulatory system for coordinated competence development and bacteriocin production. J Bacteriol 192: 1844–1852.

Perry, J.A., Jones, M.B., Peterson, S.N., Cvitkovitch, D.G., and Lévesque, C.M. (2009) Peptide alarmone signalling triggers an auto-active bacteriocin necessary for genetic competence. Mol Microbiol 72: 905–917.

Qi, F., Merritt, J., Lux, R., and Shi, W. (2004) Inactivation of the *ciaH* Gene in *Streptococcus mutans* diminishes mutacin production and competence development, alters sucrose-dependent biofilm formation, and reduces stress tolerance. Infect Immun 72: 4895–4899.

Reck, M., Tomasch, J., and Wagner-Döbler, I. (2015) The Alternative Sigma Factor SigX Controls Bacteriocin Synthesis and Competence, the Two Quorum Sensing Regulated Traits in *Streptococcus mutans*. PLOS Genetics 11: e1005353.

Ricomini, F., Antonio, P., Khan, R., Åmdal, H.A., and Petersen, F.C. (2019) Conserved pheromone production, response and degradation by *Streptococcus mutans*. bioRxiv635508.

Senadheera, D.B., Cordova, M., Ayala, E.A., Chavez de Paz, L.E., Singh, K., Downey, J.S., et al. (2012) Regulation of bacteriocin production and cell death by the VicRK signaling system in *Streptococcus mutans*. J Bacteriol 194: 1307–1316.

Senadheera, M.D., Lee, A.W., Hung, D.C., Spatafora, G.A., Goodman, S.D., and Cvitkovitch, D.G. (2007) The *Streptococcus mutans vicX* gene product modulates *gtfB/C* expression, biofilm formation, genetic competence, and oxidative stress tolerance. J Bacteriol 189: 1451–1458.

Senadheera, M.D., Guggenheim, B., Spatafora, G.A., Huang, Y.C., Choi, J., Hung, D.C.I., et al. (2005) A VicRK Signal Transduction System in *Streptococcus mutans* Affects *gtfBCD*, *gbpB*, and *ftf* Expression, Biofilm Formation, and Genetic Competence Development. J Bacteriol 187: 4064–4076.

Shanker, E., and Federle, J.M. (2017) Quorum Sensing Regulation of Competence and Bacteriocins in *Streptococcus pneumoniae* and *mutans*. Genes 8: 15.

Shields, R.C., O’Brien, G., Maricic, N., Kesterson, A., Grace, M., Hagen, S.J., and Burne, R.A. (2017) Genome-wide screens reveal new gene products that influence genetic competence in *Streptococcus mutans*. J Bacteriol 200: e00508–17.

Son, M., Ahn, S., Guo, Q., Burne, R.A., and Hagen, S.J. (2012) Microfluidic study of competence regulation in *Streptococcus mutans*: environmental inputs modulate bimodal and unimodal expression of *comX*. Mol Microbiol 86: 258–272.

Son, M., Ghoreishi, D., Ahn, S.J., Burne, R.A., and Hagen, S.J. (2015) Sharply Tuned pH Response of Genetic Competence Regulation in *Streptococcus mutans*: a Microfluidic Study of the Environmental Sensitivity of *comX*. Appl Environ Microbiol 81: 5622–5631.

Son, M., Shields, R.C., Ahn, S., Burne, R.A., and Hagen, S.J. (2015) Bidirectional signaling in the competence regulatory pathway of *Streptococcus mutans*. FEMS Microbiol Lett 362: fnv159–fnv159.

Taniguchi, Y., Choi, P.J., Li, G., Chen, H., Babu, M., Hearn, J., et al. (2010) Quantifying *E. coli* Proteome and Transcriptome with Single-Molecule Sensitivity in Single Cells. Science 329: 533.

Terleckyj, B., Willett, N.P., and Shockman, G.D. (1975) Growth of several cariogenic strains of oral streptococci in a chemically defined medium. Infect Immun 11: 649–655.

Tremblay, Y.D.N., Lo, H., Li, Y., Halperin, S.A., and Lee, S.F. (2009) Expression of the *Streptococcus mutans* essential two-component regulatory system VicRK is pH and growth-phase dependent and controlled by the LiaFSR three-component regulatory system. Microbiology 155: 2856–2865.

Underhill, S.A.M., Shields, R.C., Kaspar, J.R., Haider, M., Burne, R.A., and Hagen, S.J. (2018) Intracellular Signaling by the *comRS* System in *Streptococcus mutans* Genetic Competence. mSphere 3: e00444–18.

Wilkening, R.V., Chang, J.C., and Federle, M.J. (2016) PepO, a CovRS-controlled endopeptidase, disrupts *Streptococcus pyogenes* quorum sensing. Molecular Microbiology 99: 71–87.

