## Supplemental Information for "Carbohydrate and PepO control bimodality in competence development by *Streptococcus mutans*"

& Stephen J. Hagen

- Figure S1: Growth of wild type and *ΔtreR* strains in different carbohydrates
- Figure S2: *cipB* is required for CSP-induction of *comX* in trehalose & maltose
- Figure S3: Growth of UA159 in buffered TV medium
- Table S1: Hill function fit parameters for CSP single cell experiments in TV

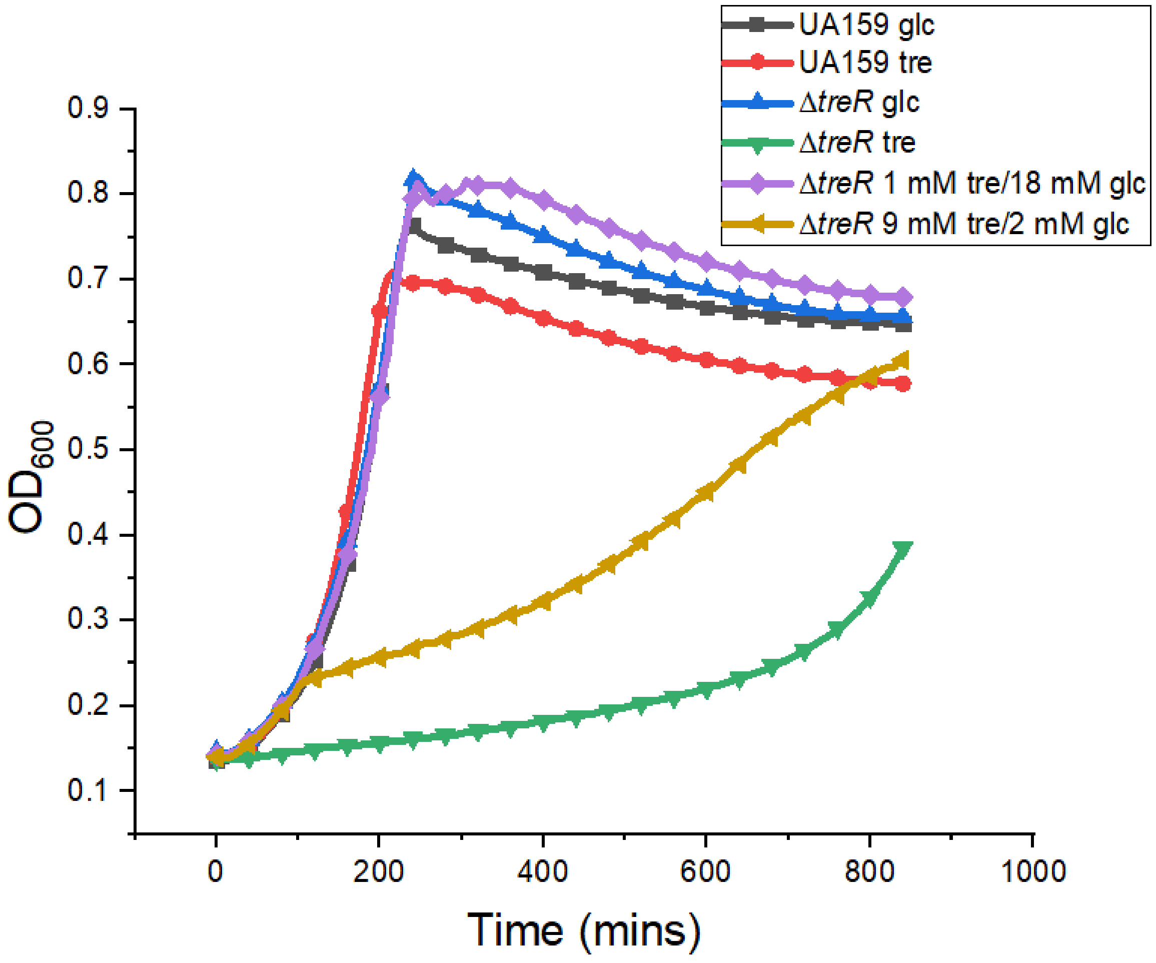


**Figure S1 – Growth of wild type and Δ*treR* strains in different carbohydrates**

Growth curves of cells with indicated genetic backgrounds in 20 mM glucose, 10 mM trehalose or a mixture of glucose and trehalose subject to the constraint 2[tre] + [glc] = 20 mM. The Δ*treR* strain clearly grows poorly on trehalose compared to the wild type, with normal growth in glucose.

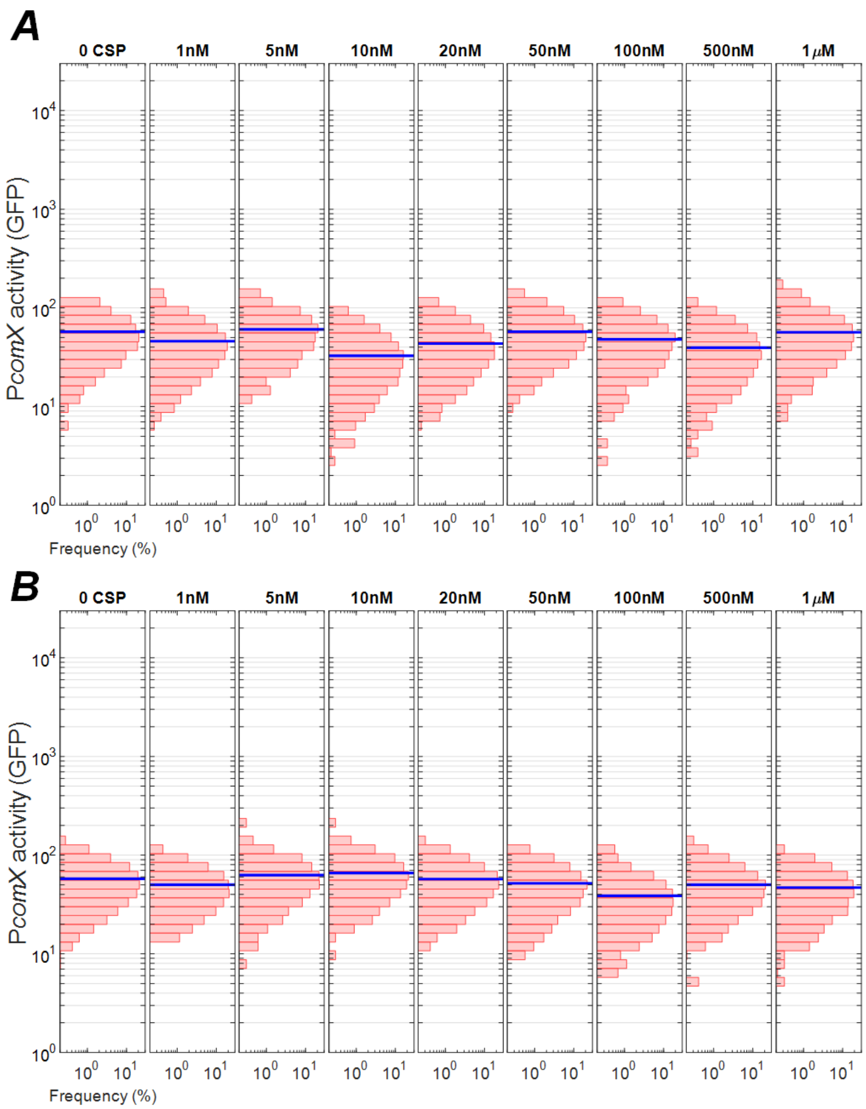


**Figure S2: *cipB* is required for CSP-induction of *comX* in trehalose & maltose**

P*comX-gfp* Δ*cipB* cells grown in TV with (A) maltose and (B) trehalose were exposed to CSP-18 at concentrations indicated at the top of each histogram for 2 hours before imaging. Histograms show individual cell fluorescence from microscopy images of cells dispersed on a glass coverslip. No *comX* activity above baseline is detected, confirming that *cipB* is required to transmit the CSP signal to ComRS in all carbohydrates studied.

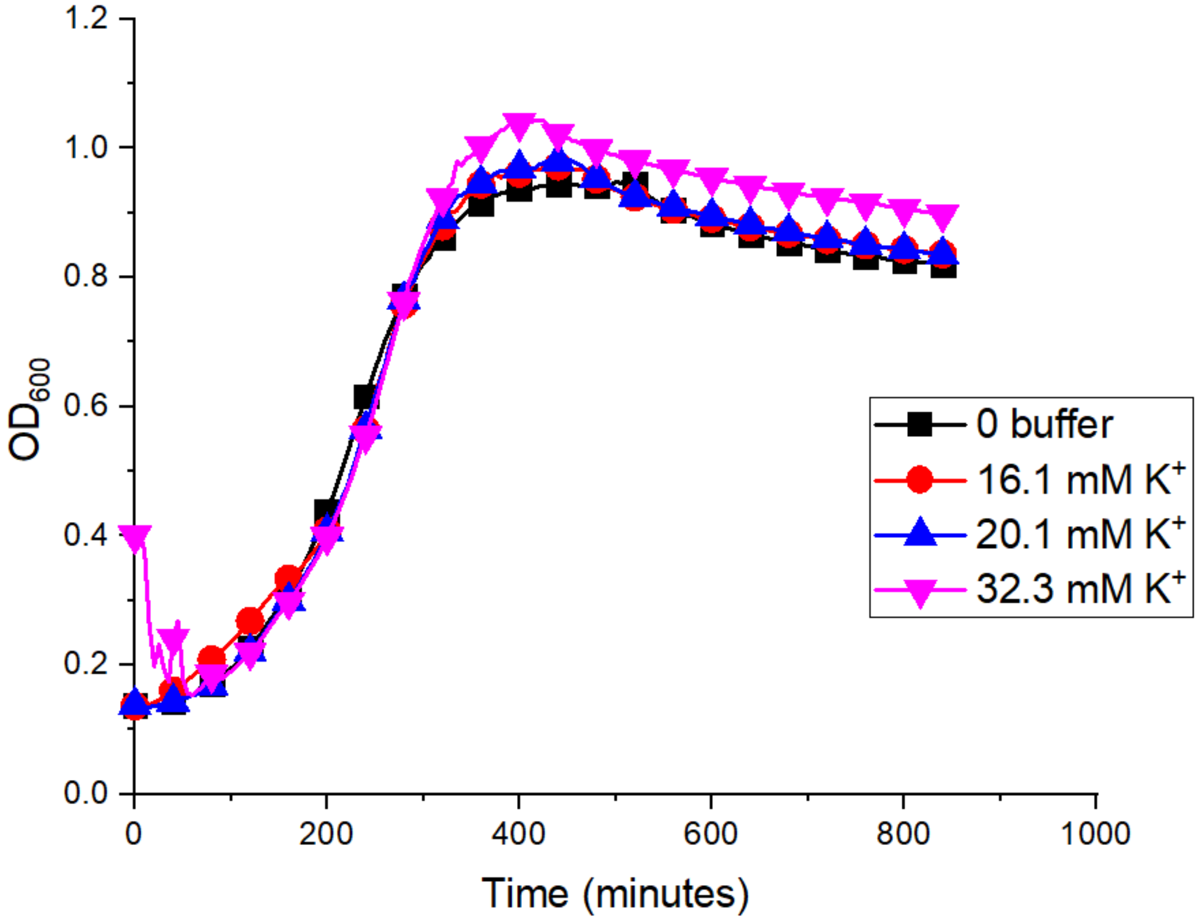


**Figure S3: Growth of UA159 in buffered TV medium**

Growth curves of cells grown in TV medium supplied with phosphate buffer diluted into the medium at the indicated potassium concentration. No effect of the buffer added to the TV is evident when comparing the black curve (no buffer) to the others.

| **Strain/sugar** | ***K* (nM)** | **g (max. % responding)** |
|---|---|---|
| Wild type -glucose | 16 ± 12 | 22 ± 9 |
| Wild type –trehalose | 7 ± 2 | 47 ± 3 |
| Wild type – maltose | 10 ± 4 | 38 ± 5 |
| Δ*pepO* - glucose | 4 ± 1 | 91 ± 0.5 |
| Δ*pepO* - trehalose | 2 ± 0.5 | 97 ± 0.3 |
| Δ*pepO* – maltose | 2 ± 1 | 96 ± 1 |
| Δ*pepO* $pepO^{+}$ - glucose | 11 ± 3 | 32 ± 3 |

**Table S1: Hill function fit parameters for CSP single cell experiments in TV**

Proportion *P* of cells activating *comX* in response to CSP was fit to

$P = g$ [CSP]/([CSP]+$K$) for indicated strains growing in TV with different carbohydrates. Parameters *K* and *g* are given.
